# The Musashi proteins are post-transcriptional activators of protein expression and alternative exon splicing in vertebrate photoreceptors

**DOI:** 10.1101/2022.03.09.483688

**Authors:** Fatimah Matalkah, Bohye Jeong, Macie Sheridan, Eric Horstick, Visvanathan Ramamurthy, Peter Stoilov

## Abstract

The Musashi proteins, MSI1 and MSI2, are conserved RNA binding proteins with a role in the maintenance and renewal of stem cells. Contrasting with this role, retina and terminally differentiated photoreceptor cells express high levels of MSI1 and MSI2, indicating that the two proteins have a role unrelated to maintaining undifferentiated cell state. Here we show that the Musashi proteins are essential in mature photoreceptors. Combined knockout of *Msi1* and *Msi2* lead to loss of the retina response to light and progressive photoreceptor cell death. The two proteins are fully redundant as individual deletion of *Msi1* or *Msi2* did not produce a phenotype. To define the molecular functions underlying the requirement for Musashi in photoreceptors, we delineated their RNA targets and analyzed the effect of the combined *Msi1*/*Msi2* knockout on transcript levels, pre-mRNA splicing, and protein expressions. We show distinct nuclear and cytoplasmic functions for the Musashi proteins in photoreceptor cells. In the nucleus, Musashi binding to the downstream proximal intron promotes splicing of alternative exons. Surprisingly, four conserved photoreceptor-specific alternative exons in genes critical for vision proved to be dispensable, leaving open questions about the selective pressures that lead to the conservation of these exons and the contribution of alternative splicing to the phenotype of the Musashi knockout. In the photoreceptor cell cytoplasm MSI1 and MSI2 act as activators of protein expression. The combined knockout of *Msi1* and *Msi2* reduced the levels of multiple proteins including proteins required for vision and photoreceptor survival.

## INTRODUCTION

Mammals express approximately 500 RNA binding proteins that associate with mRNA and pre-mRNA. These RNA binding proteins regulate pre-mRNA processing, mRNA localization, mRNA stability, and translation (Gerstberger et al., 2014). Networks of RNA binding proteins, specific to, or highly expressed in neurons perform critical roles that range from diversifying the transcriptome through alternative splicing and poly-adenylation to directing transport of specific mRNA to cellular compartments for localized translation (Agrawal and Welshhans, 2021; Darnell and Richter, 2012; Furlanis and Scheiffele, 2018; Gerstberger et al., 2014). Neuronal RNA binding proteins are essential for the development and function of the central nervous system. Processes such as axon guidance, synaptic plasticity, cell survival and cell excitation are reliant on the activity of RNA binding proteins (Furlanis and Scheiffele, 2018). Many RNA binding proteins belong to families of orthologs that share varying degrees of sequence homology and functional redundancy. Interestingly, even orthologous proteins with highly similar sequences and biochemical properties are not fully redundant and can have distinct roles in the nervous system. This divergence in function among orthologues can be derived from differences in expression levels across different cell types, subcellular localization, or their interactomes (Iijima et al., 2014; Lee et al., 2016, 2009; Saito et al., n.d.).

Neurons stand out among other cell types by the pervasive use of alternative exons (Irimia et al., 2014; Li et al., 2014). The wide use of alternatively spliced exons in neurons is due to the absence of a major splicing repressor, PTBP1, and the expression of neuron specific splicing factors (Furlanis and Scheiffele, 2018). While the importance of neuronal splicing programs is well established through knockouts of splicing regulators, the functions of the individual alternative exons are less clear. Functional significance of individual exons is commonly assigned based on sequence conservation and the nature of the protein hosting the exon. More recently, an empirical picture of the functional impact of individual exons has started to emerge. Functions of individual exons range from fine-tuning protein interactions to regulatory switches that shut down protein expression (Gonatopoulos-Pournatzis and Blencowe, 2020; Lin et al., 2020; Möröy and Heyd, 2007). Interestingly, in several cases, deletion of conserved alternative exons has failed to produce an obvious phenotype in mouse models *in vivo* (Fukuda et al., 2002; Gimond et al., 1998; Homanics et al., 1999; Vrhovski et al., 2004). The existence of exons that appear to be dispensable raises the question if the isoforms containing these exons are important under conditions that are not typically tested, or if such exons are neutral sequences that do not affect the function of the protein.

Among the RNA binding proteins expressed in neurons are the Musashi family of proteins MSI1 and MSI2 (Okano, 2002; Sakakibara et al., 1996; Sakakibara and Okano, 1997). The Musashi proteins bind to UAG sequences on RNA through two RNA recognition motifs at their N-terminus (Iwaoka et al., 2017; Lan et al., 2020). The founding member of the family, the *Drosophila Msi* protein was first described as a factor that maintains the undifferentiated state of stem cells by repressing the translation of the *Notch* regulator *Numb* (Imai et al., 2001). This function of Musashi is preserved in vertebrates where its homologues, MSI1 and MSI2, are required for stem cell maintenance and are investigated for their role in cancer progression (Fox et al., 2015; Kharas et al., 2010; Li et al., 2015; Sakakibara et al., 2002). Subsequent studies showed that the effect of the Musashi proteins on translation is context and substrate dependent, and they can positively or negatively regulate protein translation by binding to the 3’-UTR of their target mRNAs (Charlesworth et al., 2006; Cragle and MacNicol, 2014; Cragle et al., 2019; Kawahara et al., 2008).

Recently we showed that in photoreceptor cells the Musashi proteins regulate alternative splicing to produce highly photoreceptor-specific isoforms of ubiquitously expressed proteins (Murphy et al., 2016; Sundar et al., 2020). The Musashi proteins maintain an exceptionally high protein level in the mature retina and their expression is developmentally regulated (Sundar et al., 2020). MSI1 levels rise sharply after birth, peak between postnatal days 2 and 4, and decline afterwards. Concomitant with the decline of MSI1 protein levels, the levels of MSI2 increase and remain constant in adulthood (Sundar et al., 2020). Single deletion of *Msi1* or *Msi2* in committed rod photoreceptor progenitor cells showed that the two paralogs are partially redundant and appear to act at time-points of retinal development that correlate with the pattern of their expression (Sundar et al., 2020). The single deletion of *Msi1* resulted in an early visual defect that was observed at the time of eye-opening in mice (postnatal day 16). In contrast, the removal of *Msi2* resulted in normal vision at postnatal day 16 that progressively declined with age (Sundar et al., 2020). Based on these findings, we proposed that MSI2 is involved in the maintenance of mature photoreceptor cells, while MSI1 functions mostly in photoreceptor precursors and immature photoreceptor cells.

To understand the role of Musashi proteins in mature photoreceptor cells, we utilized an inducible mouse model to delete the Musashi genes in mature photoreceptors. We find the Musashi proteins to be essentials for the function and viability of the photoreceptors. To our surprise, despite the reciprocal regulation of MSI1 and MSI2 levels during development, the two proteins are fully redundant in mature photoreceptor cells. To understand the molecular mechanisms underlying the function of the Musashi proteins in photoreceptors, we identified the transcripts recognized by MSI1 and MSI2 *in vivo* and investigated how deletion of the *Msi1* and *Msi2* genes affected pre-mRNA splicing, global transcript levels, and protein expression. We demonstrate *in vivo* that the Musashi proteins bind downstream of photoreceptor-specific exons to activate their splicing. In addition, we show that in photoreceptors the Musashi proteins act almost exclusively as post-transcriptional activators of protein expression.

## RESULTS

### Induced depletion of *Msi1* and *Msi2* in mature photoreceptor cells

We recently showed that the Musashi proteins are expressed at exceptionally high levels in the retina (Sundar et al., 2020). The two proteins show distinct developmental regulation in the retina after birth that correlated with the phenotypes of *Msi1* and *Msi2* in photoreceptor precursors cells (Sundar et al., 2020). This correlation suggested separate roles for MSI1 and MSI2 in developing and mature photoreceptor cells, respectively. We tested the roles of the Musashi protein in mature photoreceptors by using tamoxifen-inducible *Cre^ERT2^* under the control of rod-specific *Pde6g* promoter to remove *Msi1* and *Msi2* in mature rod-photoreceptor cells (Koch et al., 2015). Floxed (*Msi1^flox/flox^*/*Msi2^flox/flox^*) mice hemizygous for *Pde6g-Cre^ERT2^* were injected with tamoxifen for three consecutive days starting at postnatal day 30 to create combined *Msi1*/*Msi2* knockout mice. Littermates carrying the floxed alleles for *Msi1* and *Msi2* (*Msi1^flox/flox^*/*Msi2^flox/flox^*) and treated with tamoxifen were used as controls. We will refer to the *Msi1^flox/flox^*/*Msi2^flox/flox^*–treated with tamoxifen as *Msi1^+/+^/Msi2^+/+^* and the mice with *Msi1^flox/flox^*/*Msi2^flox/flox^-Pde6gCre^ERT2^* treated with tamoxifen as *Msi1^-/-^*/*Msi2^-/-^*. Immunocytochemistry (ICC) demonstrated that, 14 days after the first tamoxifen injection, the MSI1 and MSI2 proteins were depleted specifically from the photoreceptors (Figure 1A). Consistent with the ICC data, western blot analysis showed two fold decrease in the MSI1 and MSI2 protein levels in retinal lysates from the *Msi1^-/-^*/*Msi2^-/-^* mice (Figure 1B and Figure 7 - figure supplement 1).

**Figure 1.**
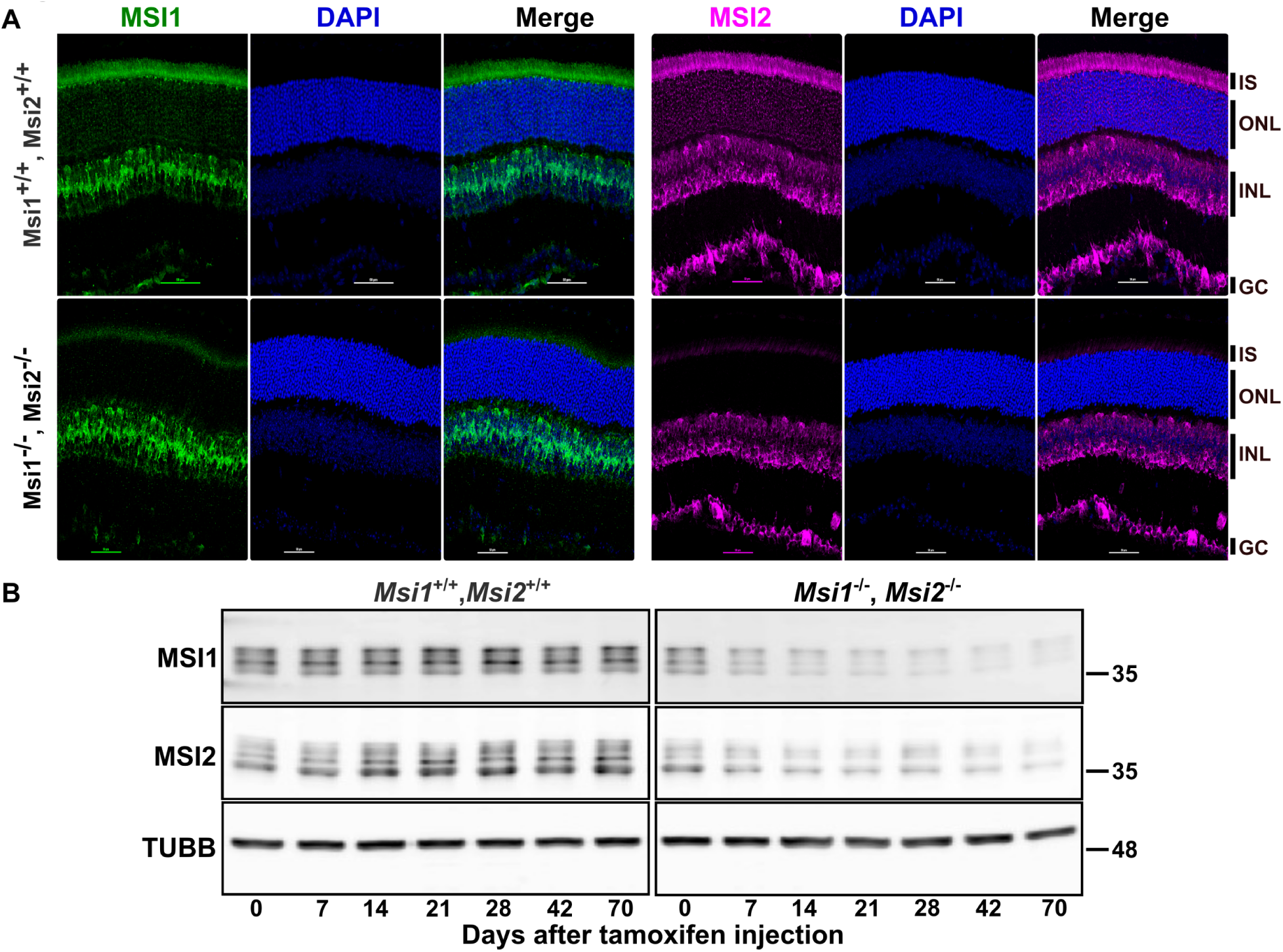
Induced double knockout of *Msi1* and *Msi2* in photoreceptor cells. **A)** Immunofluorescence micrographs of retinal cross-sections collected at day 14 post tamoxifen injection from *Msi1*^+/+^/*Msi2*^+/+^ littermate control and *Msi1*^-/-^/*Msi2*^-/-^, stained for MSI1(green), MSI2 (magenta), and DAPI (blue). ONL: outer nuclear layer (photoreceptor cell layer). INL: inner nuclear layer. GC: ganglion cell layer. Objective, 40x. **B)** Immunoblot of lysates from *Msi1*^+/+^/*Msi2* ^+/+^ and *Msi1*^-/-^ /*Msi2*^-/-^ retinas collected between 0 and 70 days after tamoxifen injections and probed with antibodies to MSI1, MSI2 and TUBB (β-tubulin, loading control).

### MSI1 and MSI2 are required for the function and survival of mature photoreceptors

To evaluate the functional significance of Musashi proteins in mature photoreceptors, we used electroretinograms (ERG) to measure the retinal response to light. We measured the dark-adapted (scotopic) and the light-adapted (photopic) responses that reflect rod and cone photoreceptor function, respectively (Benchorin et al., 2017). We used repeated measures two-way ANOVA to determine the effect of the genotype and time post injection on the ERG A-Wave amplitude. Two-way repeated measures ANOVA found a significant interaction of the genotype and the time after injection (Scotopic response: F(12,1)= 19.47, p-value<0.0001; Photopic response: F(12,1)=10.37, p-value<0.0001). The response to light of the *Msi1^-/-^/Msi2^-/-^* animals and the age-matched *Msi1^+/+^/Msi2^+/+^* controls was significantly different 28 days after tamoxifen injection (Figure 2A, B). The response to light continued to decrease rapidly thereafter and was nearly undetectable by day 105 post-injection (Figure 2A, B).

**Figure 2.**
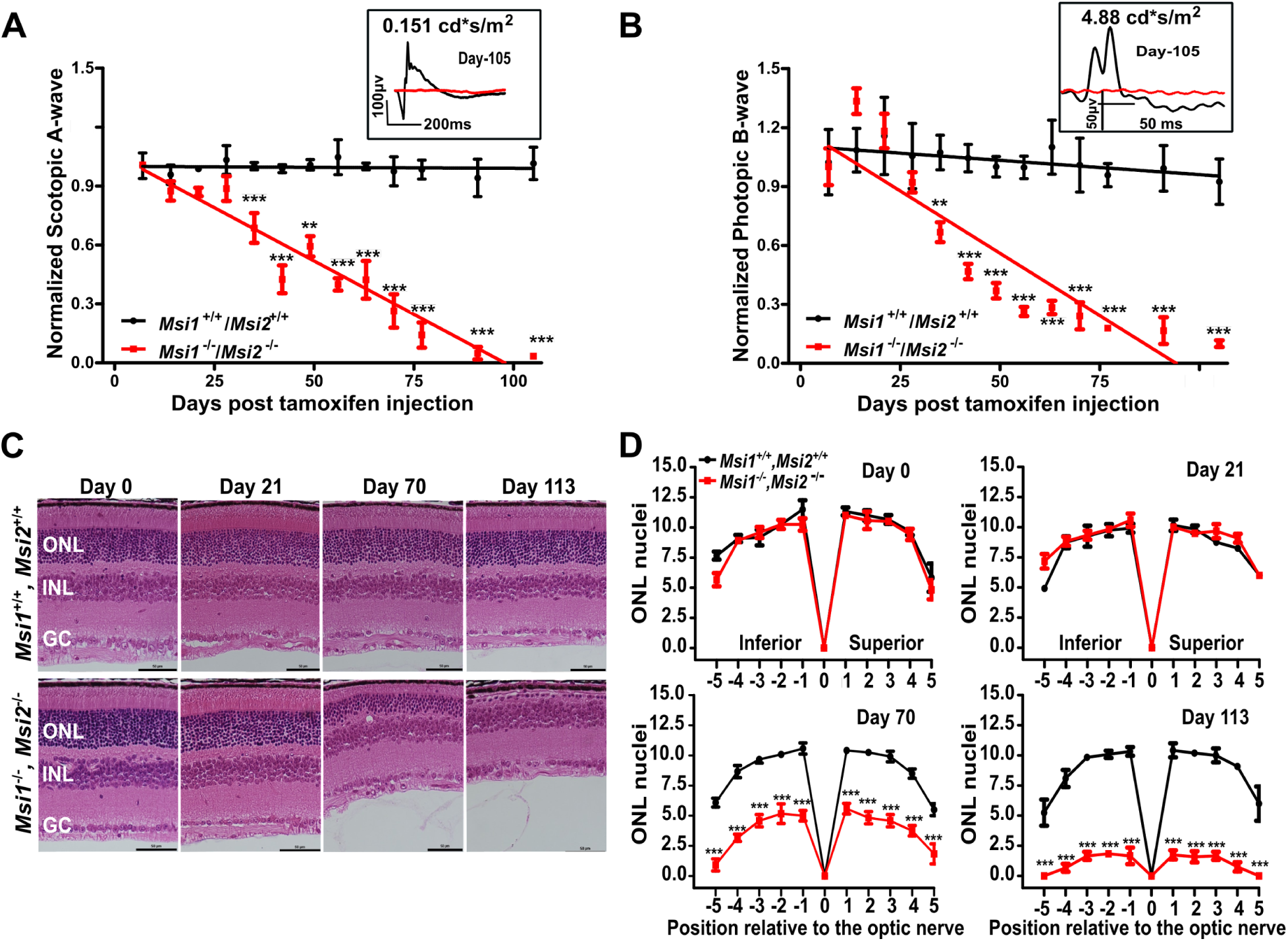
Progressive loss of response to light and retinal degeneration after double knockout of *Msi1* and *Msi2* in photoreceptor cells. Time course of scotopic A-wave (**A**) and photopic B-wave (**B**) responses from *Msi1^+/+^/Msi2 ^+/+^* (black line), and *Msi1 ^-/-^ /Msi2 ^-/-^* retinas (red line) following tamoxifen injection. Scotopic and photopic waveforms were obtained 0.151 cd*s/m^2^ and 4.88 cd*s/m^2^ flashes, respectively. Insets show representative electroretinograms from a single *Msi1^+/+^/Msi2 ^+/+^* (black line) and *Msi1 ^-/-^ /Msi2 ^-/-^* (red line) mouse retina at day 105 post tamoxifen injection. The data points from the scotopic and photopic responses are represented as mean ± SEM of 8 eyes (4 animals). Pairwise t-test with Bonferroni correction for multiple comparisons was used to determine the effect of genotype on the A-wave amplitude at each time point. Significance levels of the pairwise comparisons is indicated as: * p-value < 0.05, ** p-value < 0.01, *** p-value < 0.001. C) Outer nuclear layer degeneration in *Msi1^-/-^/Msi2^-/-^* knockout retinas. Representative H&E-stained sections from the *Msi1*^+/+^/*Msi2*^+/+^ and *Msi1^-/-^/Msi2^-/-^* retinas collected between day 0 and day 113 post-tamoxifen injection. ONL: outer nuclear layer (Photoreceptor nuclei), INL: inner nuclear layer, GC: ganglion cells. 40X objectives and scale bar represents 50 μm. D) Spider plots displaying the thickness of the ONL as the number of nuclei measured at ten points stepped by 400µm from the optical nerve at different time points post-tamoxifen injection. Data are shown as mean ± SEM. Pairwise t-test with Bonferroni correction for multiple comparisons was used to determine the effect of the genotype on the outer nuclear layer thickness at each time point. Significance levels of the pairwise comparisons is indicated as: * p-value < 0.05, ** p-value < 0.01, *** p-value < 0.001.

To assess the retinal morphology following *Msi1/Msi2* deletion, we performed hematoxylin and eosin (H&E) staining on retinal cross-sections collected from *Msi1^-/-^/Msi2^-/-^* and control mice between days 0 and 113 post-injection (Figure 2C, D, and Figure 2 - figure supplement 1). A significant effect of the genotype over time was confirmed by two-way ANOVA (F(1,259)=61.04, p-value<0.0001). In agreement with the ERG data, we did not observe significant morphological changes up to 28 days after tamoxifen injection (Figure 2C, D, and Figure 2 - figure supplement 1). After day 28 post-injection, we observed progressive degeneration of the photoreceptor cell layer (Figure 2D, Figure 2 - figure supplement 1). Approximately half of the photoreceptor cells were lost by day 42 post-injection in the knockout retinas, with a single layer of photoreceptor cells remaining at day 113 post-injection (Figure 2 C, D, and Figure 2 - figure supplement 1). We did not observe any significant changes in the inner retina, including the inner nuclear layer (INL) and the ganglion cell layer (GCL). Our results show that the combined deletion of *Msi1/Msi2* in mature photoreceptors leads to a rapid and progressive decline in the function and viability of photoreceptor cells that starts three to four weeks after depletion of the Musashi proteins.

### MSI1 and MSI2 are redundant in the maintenance of mature photoreceptors

To determine if MSI2 plays a dominant role in mature retina, as the developmental regulation of MSI1 and MSI2 protein expression would suggest, we used the tamoxifen-inducible *Cre-LoxP* system described above to delete the *Msi1* and *Msi2* genes individually in mature photoreceptors. We confirmed the photoreceptor-specific loss of MSI1 and MSI2 protein by immunostaining retinal cross-sections obtained at day 14 post-injection (Figure 3A, C). Western blot analysis of retinal lysates showed that tamoxifen injection required 7 to14 days to ablate the proteins, in agreement with our observation of the double knockout (Figure 3B, C). The MSI2 protein level was upregulated 1.4-fold in the *Msi1^-/-^* retina compared to the control; however, the increase did not reach statistical significance at the number of replicates used (n=3).

**Figure 3.**
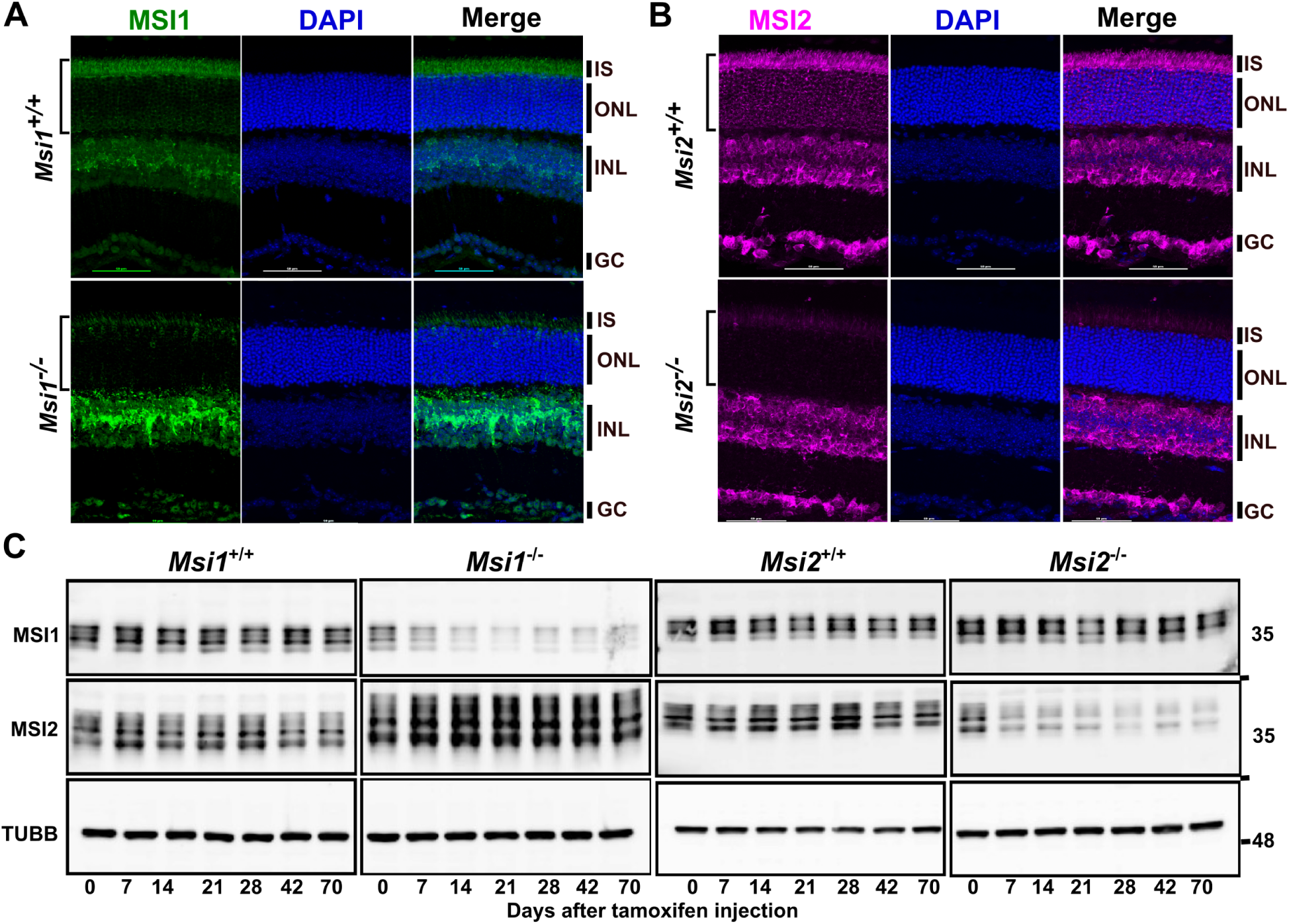
Induced single knockouts of *Msi1* or *Msi2* in photoreceptor cells. **A)** Immunofluorescence micrographs of retinal cross-sections collected at D14 post-tamoxifen injection from *Msi1*^-/-^ mice and *Msi1*^+/+^ littermates **(A)** or *Msi2*^-/-^ mice and *Msi2*^+/+^ littermates **(B).** Sections were stained with antibodies to MSI1(green), MSI2(magenta) and DAPI (blue). ONL: outer nuclear layer (photoreceptor cell layer). INL: inner nuclear layer. GC: ganglion cell layer. Objective,40X. **C)** Immunoblot of lysates from *Msi1*^+/+^ and *Msi1*^-/-^ retinas collected between 0 and 70 days after tamoxifen injections and probed with antibodies to MSI1, MSI2 and TUBB (loading control).

Neither *Msi1* nor *Msi2* single ablation had an effect on retina function (Figure 4A). The scotopic and photopic ERG responses collected from day 7 to day 230 post injection show that the response to light of the single *Msi1* or *Msi2* knockout mice are indistinguishable from the control animals (Figure 4A, and Figure 4 - figure supplement 1A, B). Similarly, the histology of the knockout retinas collected at day 230 post injection do not show signs of degeneration (Figure 4B, C). Together, these data demonstrate that Musashi proteins are fully redundant in mature photoreceptors.

**Figure 4.**
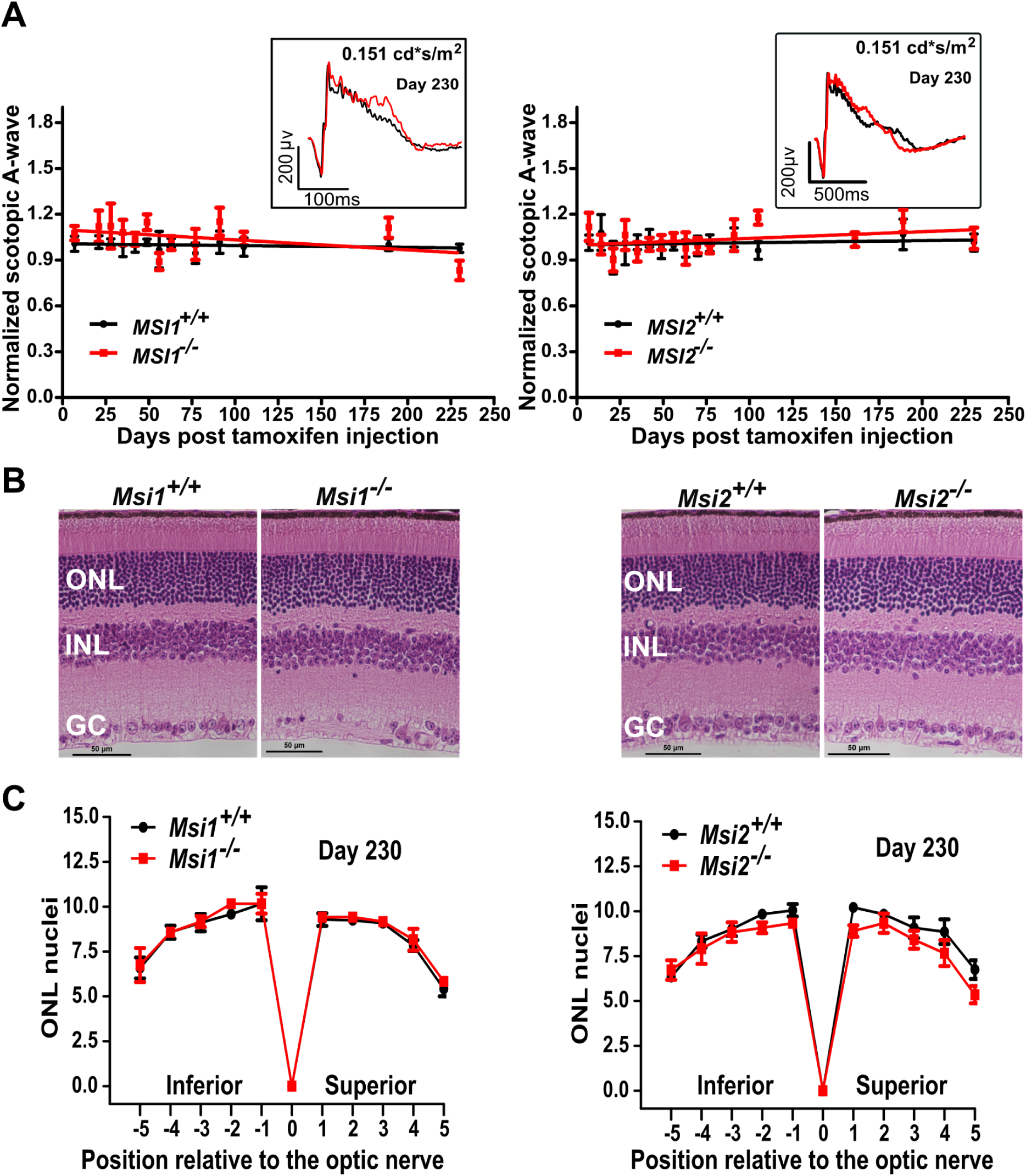
Normal photoreceptor response to light and retinal morphology in the single knockouts *Msi1* or *Msi2* in photoreceptor cells. **A)** Scotopic mean A-wave response from single *Msi1 ^-/-^* (left) or *Msi2 ^-/-^* (right) knockouts in photoreceptors (red lines) and littermate controls (black lines) recorded between 0 and 230 days days post tamoxifen injection. Scotopic waveforms were obtained using 0.151 cd-s/m^2^ flashes. The insets of panels A and B show representative scotopic (dark-adapted) electroretinograms from a single knockout (red) or control (black) retina at D230 post-injection using 0.151 cd-s/m^2^ flashes. Scotopic waveforms were obtained after overnight dark adaptation using 0.151 cd-s/m^2^ flashes. **B)** Representative H&E-stained sections from retinas of single *Msi1 ^-/-^* (left) or *Msi2 ^-/-^* (right) knockouts in photoreceptor cells and littermate controls collected 230 days post-tamoxifen injection. ONL: outer nuclear layer (Photoreceptor nuclei), INL: inner nuclear layer, GC: ganglion cells. 40X objectives, scale bar, represents 10μm. **C)** Spider plots of ONL thickness for single *Msi1 ^-/-^* (left) or *Msi2 ^-/-^* (right) knockouts (red) and littermate controls (black). The plots display the thickness of the ONL as the number of nuclei measured at ten points stepped by 400µm from the optical nerve in retinas collected 230 days after tamoxifen injection. Data are shown as mean ± SEM of 8 eyes, from 4 animals.

### Binding of Musashi downstream of alternative exons promotes their inclusion in photoreceptor cells

To delineate the transcripts bound by Musashi and the positions on the targets where Musashi binds, we used UV Cross-Linking and Immuno-Precipitation followed by high throughput sequencing of the associated RNA fragments (eCLIP-Seq). The MSI1 and MSI2 proteins are fully redundant in the adult retina, their RNA binding domains are 77% (RRM1) to 92% (RRM2) identical, and the two proteins recognize the same UAG binding site (Lan et al., 2020; Zearfoss et al., 2014). Thus, we argued that performing the eCLIP-Seq experiment on MSI1 will be sufficient to identify the targets for both proteins.

The eCLIP-Seq data shows that MSI1 binds predominantly to the 3’-UTRs of mRNA (59.7% of binding sites, Figure 5A, C) and introns of pre-mRNA (32.7% of binding sites, Figure 5A, E). We performed motif enrichment analysis of the sequence surrounding the eCLIP crosslink sites using the HOMER and MEME software suites. Both software packages identified as enriched motifs centered on a UAG core (Figure 5 - figure supplement 1), in agreement with the UAG binding site sequence for the Musashi proteins derived from *in vitro* binding and structural studies (Dominguez et al., 2018; Lan et al., 2020; Zearfoss et al., 2014). The crosslink frequency peaks at position -1 relative to the top motif identified by DREME, BUAG, indicated direct binding of MSI1 (Figure 5B).

**Figure 5.**
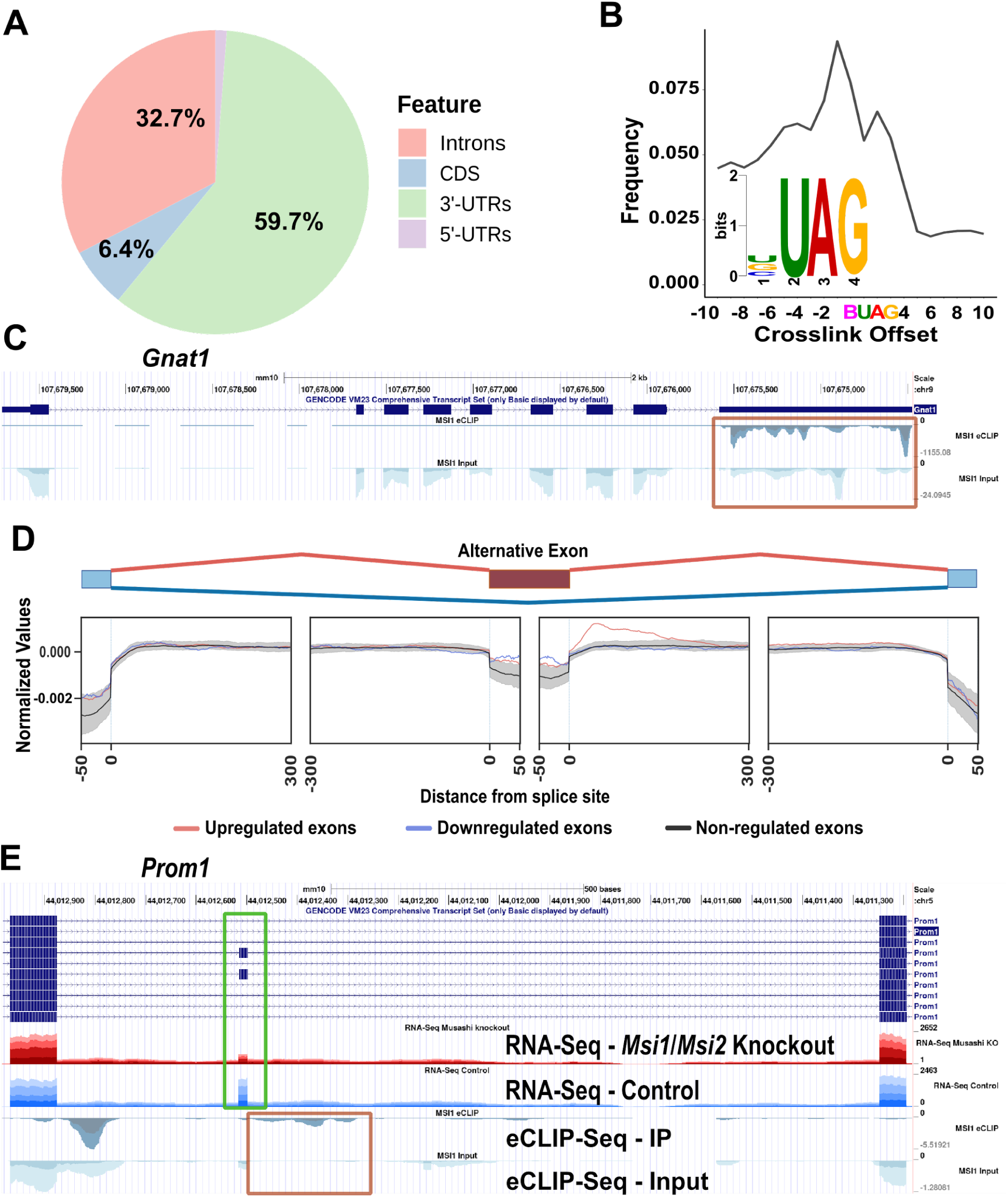
In the retina MSI1 binds to UAG motifs located predominantly in introns and 3’-UTRs. **A)** Distribution of MSI1 binding sites as identified by eCLIP-Seq on mouse retinal samples across mRNA features. **B)** eCLIP crosslink frequency relative to the top scoring motif (BUAG) identified by DREME. **C)** UCSC Genome browser tracks showing the eCLIP-seq signal enrichment over the 3’-UTR of the *Gnat1* gene (orange box). Replicates are stacked and indicated by different shading colors. Scales are 0 to -1155 for the eCLIP IP and 0 to -24 for the eCLIP input. **D)** RNA binding protein map showing MSI1 binding relative to an alternative metaexon. Exons upregulated by the Musashi proteins (downregulated in the *Msi1*/*Msi2* double knockout) are shown in red, exons downregulated by the Musashi proteins are shown in blue and alternative exons remaining unchanged in the knockout are shown in black. Gray shading indicates the 99.5% confidence interval derived from 1000 random permutations. MSI1 binding sites are enriched downstream of alternative exons upregulated by the Musashi proteins. **E)** UCSC Genome browser tracks showing MSI1 binding (orange box) downstream of an alternative exon in the *Prom1* gene regulated by the Musashi proteins (green box). RNA-Seq tracks show the read density for retinal samples derived from photoreceptor specific *Msi1*/*Msi2* double knockouts and matched controls. Replicates are stacked and indicated by different shading colors. Scales are 0 to -5.5 for the eCLIP IP and 0 to -1.3 for the eCLIP input.

Previously we identified a photoreceptor-specific alternative splicing program by comparing the splicing in *Aipl1* knockout mouse retina devoid of photoreceptor cells to wild type retina (Murphy et al., 2016). Motif enrichment analysis suggested a role for the Musashi proteins in controlling this program. Through minigene experiments and mutagenesis, we demonstrated that the splicing of at least one exon, exon 2A in the *Ttc8* gene, is activated by MSI1 bound downstream of the exon (Murphy et al., 2016). Here we sought to determine on a global scale how the Musashi proteins are regulating alternative splicing in photoreceptor cells *in vivo*. Analysis by RNA-Seq of alternative splicing in *Msi1^-/-^/Msi2^-/-^* retina 21 days after tamoxifen injection identified 165 exons that had reduced inclusion levels and 115 exons that were upregulated in the knockout (Supplementary Tables 1, 2, and 3). Out of the 165 exons downregulated in the *Msi1*/*Msi2* knockout, 52 were also significantly downregulated in the photoreceptor-devoid retina of the *Aipl1* knockout mice, with another 40 exons showing the same direction of change but not reaching statistical significance in the *Aipl1* knockout retina (Supplementary Table 2) (Murphy et al., 2016). None of the significantly downregulated exons in the *Msi1*/*Msi2* knockout retina were significantly upregulated in the *Aipl1* knockout (Supplementary Table 3). We did not observe a correlation between the exons significantly upregulated in the *Msi1^-/-^/Msi2^-/-^* retina and the exons differentially spliced in the *Aipl1* knockout retina.

Our previous work suggested that the Musashi proteins promote inclusion of alternative exons by binding downstream of the exon in the adjacent intron. To determine if this mode of regulation is common *in vivo* we combined the eCLIP-Seq and RNA-Seq data to build an RNA splicing map of a meta cassette exon (Figure 5D). The splicing map shows significant enrichment of Musashi protein binding in the downstream introns proximal to the exons upregulated by the Musashi proteins (exons downregulated in the knockout). No significant enrichment of Musashi binding sites was observed for exons repressed in the wild type animals compared to the Musashi knockouts. Our data demonstrates that *in vivo* Musashi proteins can promote inclusion of alternative exons by binding in the intron downstream of the exon.

### Photoreceptor-specific microexons in the *Ttc8*, *Cep290*, *Cc2d2a*, *Cacna2d4* and *Slc17a7* genes are dispensable

Considering the requirement of Musashi proteins for vision and their role in promoting splicing of photoreceptor specific exons, we next tested if photoreceptor-specific alternative splicing variants are required for vision. Using CRISPR/Cas 9 we deleted photoreceptor-specific exons in the *Ttc8*, *Cep290*, *Cc2d2a*, *Cacna2d4* and *Slc17a7* genes (Murphy et al., 2016). The exons in *Ttc8*, *Cep290*, *Cc2d2a* and *Cacna2d4* genes are microexons showing sequence conservation across vertebrates that is traceable to fish (Figure 6 - figure supplement 1A). We further confirmed by RT-PCR that the four alternative exons are used in zebrafish and are included at high rate in the zebrafish eye (Figure 6 - figure supplement 1B). The photoreceptor-specific exon in *Slc17a7* is confined to rodents and serves as a control representing an evolutionary novel exon that is less likely to impact the function of the host protein. Four of the exons in the *Ttc8*, *Cep290*, *Cc2d2a* and *Slc17a7* genes were downregulated in our *Msi1*/*Msi2* double knockout mice.

Deletion of each exon was confirmed by sequencing the alleles after the founders have been outcrossed (Figure 6 - supplement 2). RT-PCR from retinal samples showed the expected expression of exon skipped isoform (Figure 6A). We examined the visual function of the exon knockout mice by ERG between one and twelve months of age. We did not observe significant differences in the response to light of the exon knockout mice compared to wild type controls (Figure 6B). Similarly, H&E stained retinal sections from the exon knockout mice had normal morphology (Figure 6C). The phenotypes of the individual exon knockout mice may have been too subtle to detect on their own. Thus, we crossed the *Ttc8*, *Cep290*, and *Cc2d2a* exon knockout mice to create a homozygous triple exon knockout mouse line. As all three proteins are part of the primary cilium and are critical for cilium biogenesis, we expected the individual exon knockout phenotypes to be amplified in the combined knockout. We monitored the vision in the triple knockout animals by ERG and their retinal morphology was assessed by H&E staining. As with the single exon knockout mice we did not observe changes in the function or morphology of the retina of the triple knockout animals (Figure 6B and C).

**Figure 6.**
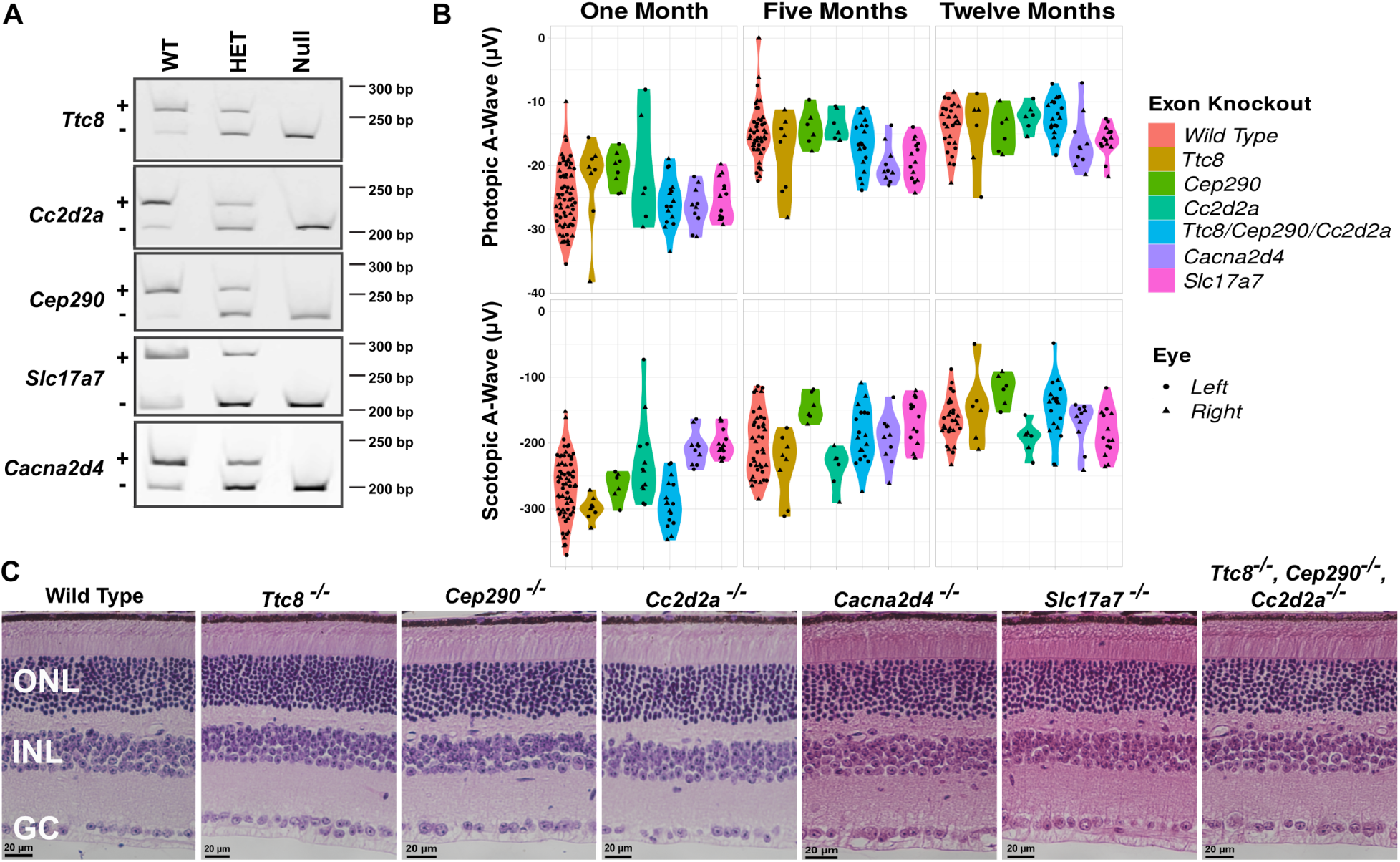
Normal photoreceptor response to light and retinal morphology of knockouts of photoreceptor specific exons in the *Ttc8*, *Cc2d2a*, *Cep290*, *Cacna2d4*, and *Slc17a7* genes. **A)** RT-PCR of retinal samples showing loss of the photoreceptor-specific mRNA isoforms in the exon knockout animals. RNA was extracted from the retinas of wild type animals (WT), and heterozygous (Het) and homozygous (Null) exon knockouts. Isoforms including and skipping the alternative exon are indicated by “+” and “-”, respectively. **B.** Violin plots of the photopic and scotopic A-wave intensities at postnatal days 30, 150 and 356 collected from the single and triple (*Ttc8*, *Cc2d2a*, *Cep290*) exon knockouts. **C)** Representative H&E-stained retinal sections from wild type mouse, knockouts of photoreceptor specific microexons in the *Ttc8*, *Cep290*, *Cc2d2a*, *Canca2d4* and *Slc17a7* genes, and combined deletion of the microexons in the *Ttc8*, *Cep290*, *Cc2d2a* genes. The samples were collected at 12 months of age. ONL: outer nuclear layer (Photoreceptor nuclei), INL: inner nuclear layer, GC: ganglion cells. 40X objectives, scale bar, represents 20μm.

### MSI1 and MSI2 are post-transcriptional activators of protein expression in photoreceptor cells

Consistent with the previously described role of the Musashi proteins in regulating mRNA translation, our CLIP-seq data showed pervasive binding of MSI1 to 3’-UTRs (Figure 5A and C). To determine the effect of Musashi on the protein expression from photoreceptor-specific transcripts, we analyzed the changes in mRNA and protein levels in *Msi1^-/-^*/*Msi2^-/-^* retina where both MSI1 and MSI2 were depleted from mature photoreceptor cells. For RNA-Seq and quantitative proteomics, we used retinas that were collected 21 days after tamoxifen injection. At this time the Musashi proteins were depleted from photoreceptor cells, while the knockout retina had normal morphology and response to light (Figure 2). Thus, the effect of mRNA and protein expression could be analyzed without the confounding effects of photoreceptor cell death.

We used isobaric labeling and mass spectroscopy to compare protein expression between knockout and control retina. We identified and quantified peptides from 8021 proteins. Of these proteins 165 showed significant differences in expression (at least 1.5 fold change in protein levels with adjusted p-value at or below 0.01) between the control and knockout retina. Of the proteins with significant changes 98 had reduced expression and 67 had elevated expression in the knockout retina (Supplementary Table 4). As expected, MSI1 and MSI2 protein levels were decreased by more than 2-fold in the retina of the *Msi1/Msi2* knockout mice (Supplementary Table 4 and Supplementary Figure 6), consistent with the change in expression observed by western blot (Figures 1, 3 and Figure 7 - figure supplement 1). Gene Ontology and KEGG pathway enrichment analysis showed that proteins downregulated in the knockout retina were strongly associated with categories related to phototransduction, photoreceptor cell structure, and photoreceptor homeostasis (Figure 7, Figure 7 - figure supplement 2, and Supplementary Table 5). The reduced expression of proteins in these categories is not a consequence of photoreceptor cell death or degeneration for two reasons. First, morphologically and physiologically, the retina of the knockout animals appeared normal at this stage. More importantly, the levels of multiple proteins that are photoreceptor-specific or are abundantly expressed in photoreceptor cells were unchanged or even increased (Figure 7). Examples include proteins with functions in phototransduction (RCVRN), outer segment structure (PRPH2, PROM1), primary cilium structure (CC2D2A, CEP290), intraflagellar transport (IFT80, IFT140), ion transport (ATP1B2), and protein transport (RD3).

**Figure 7.**
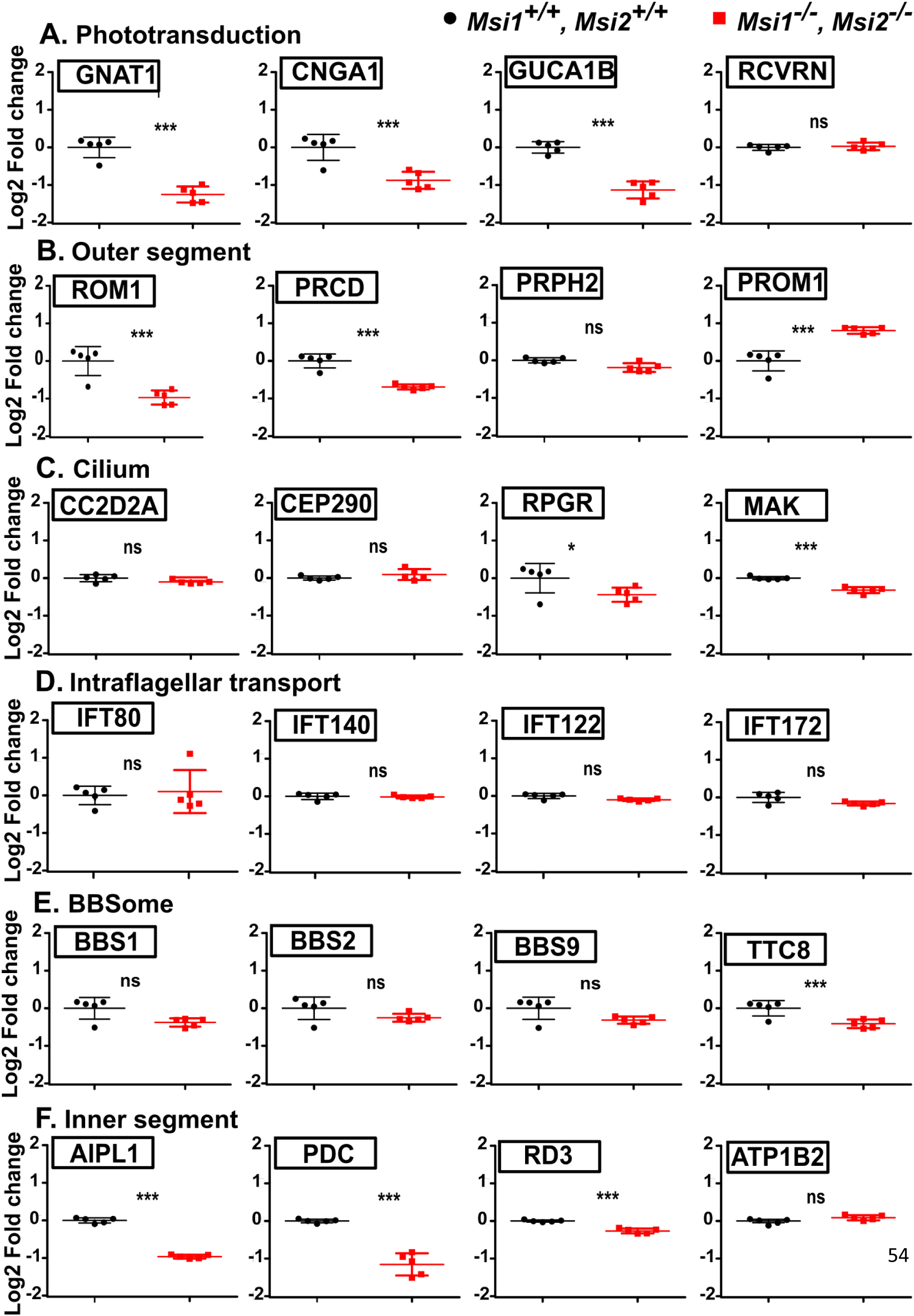
Expression of proteins critical for photoreceptor function after induced knockout of *Msi1* and *Msi2* in photoreceptor cells. Boxplots representing the log2 of the fold change in protein expression relative to the median of the control. Retinal samples were collected 21 days after inducing the knockout at postnatal day 30. Protein levels were determined using isobaric labeling and mass-spectroscopy. Boxplots represent a selected set of proteins that are components of the phototransduction pathway (**A)**, outer segment structure **(B)**, primary cilium structure **(C)**, intraflagellar transport complex **(D)**, BBSome **(E)**, and the inner segment **(F)**. Significance level is indicated as: * adjusted p-value < 0.05, ** adjusted p-value < 0.01, *** adjusted p-value < 0.001.

In contrast to the downregulated proteins, most of the proteins with increased expression in the knockout retina were associated with Gene Ontology terms and KEGG pathways involved in cell proliferation, extracellular matrix structure, immune response, and angiogenesis (Supplementary Table 5). Closer examination of the upregulated proteins revealed proteins (GFAP, CLU, STAT3, JUNB, IRF9, A2M, B2M, complement components) that are expressed at elevated levels across various models of retinal degeneration (Bertrand et al., 2021, p. 27, 2021, p. 27; Cheng and Molday, 2013; Hackam et al., 2004; Uren et al., 2014). Single cell RNA-Seq of the Cwc27^fs^ model of retinal degeneration further indicated that many of the upregulated genes are expressed by glia (Bertrand et al., 2021). To determine how MSI1 and MSI2 regulate protein expression in photoreceptor cells, we defined two sets of genes. The first set were genes that are either specifically expressed in photoreceptor cells or have at least two fold higher expression in photoreceptors compared to any other cell type in the retina. The second set contained genes that are either not expressed in photoreceptor cells or have at least two fold lower expression compared to other retinal cell types. The two sets of genes were derived from differential expression analysis of single cell RNA-Sequencing data by Macosco et al (Macosko et al., 2015). We will use “photoreceptor-specific” as a shorthand for the subset of genes highly expressed specifically in photoreceptor cells, with the understanding that many of these genes are also expressed in other cell types, albeit at lower levels. Most proteins with significantly lower expression in the *Msi1^-/-^/Msi2^-/-^* retina and without a significant change in their mRNA levels, belonged to the photoreceptor-specific set of genes (Figure 8A and Supplementary Table 7). Only two photoreceptor-specific proteins, PROM1 and IMPG2, had markedly higher expression in the knockouts (Figure 8A). A cumulative plot of the changes in protein and RNA expression from the photoreceptor-specific genes shows a global trend in reduced protein levels that were not matched by a corresponding decrease in transcript levels (Figure 8B). Taken together our data demonstrates that the Musashi proteins promote protein expression post-transcriptionally.

**Figure 8.**
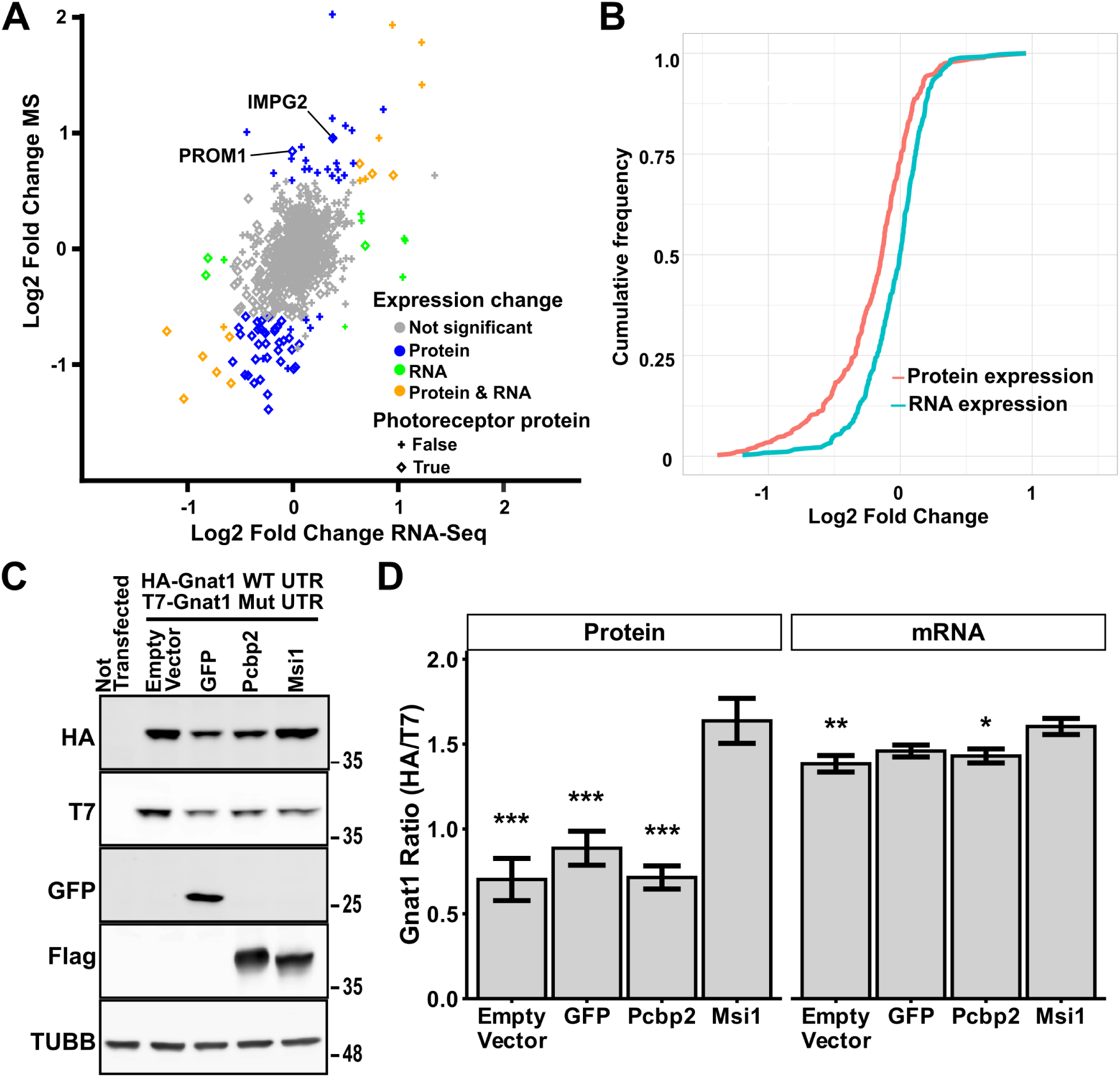
In photoreceptor cells MSI1 and MSI2 act to promote protein expression post-transcriptionally. **A)**. Scatter plot comparing protein and transcript levels changes after double *Msi1*/*Msi2* knockout in mature photoreceptor cells. Protein expression was quantified by isobaric labeling and mass spectrometry. Transcript expression levels were determined by RNA-Seq. Genes that are highly expressed specifically in photoreceptor cells are shown as rombs and genes highly expressed outside of photoreceptor cells are shown as crosses. Color indicates significant changes in protein levels alone (blue), in mRNA levels alone (green), in both protein and mRNA levels (orange), or no significant change in expression (gray). **B)** Cumulative frequency plots showing the deletion of *Msi1/Msi2* in mature photoreceptors leads to broad decrease of photoreceptor protein levels in excess to changes in transcript levels (Kolmogorov-Smirnov p-value=1.1*10^-12^). **C)** Western blot analysis of recombinant GNAT1 expression in NIH 3T3 cells. Cells were transfected with equal amounts HA-tagged *Gnat1* clone with wild type 3’-UTR and T7-tagged *Gnat1* clone with mutant 3’-UTR that lacks Musashi binding sites. Each transfection included one of the following: empty pcDNA3.1(+) vector, vector expressing GFP, vector expressing flag-tagged *Pcbp2*, and vector expressing flag-tagged *Msi1*. Non transfected cells were included as a control for the specificity of the antibodies. TUBB serves as loading control. **D)** Ratios of the HA (wild type) to T7 (mutant) tagged GNAT1 proteins and ratios of the corresponding RNA transcripts in the transfected NIH 3T3 cells. The statistical significance of the effect of MSI1 in pairwise comparisons to the controls was determined by Tukey HSD. Significance level is indicated as:* p-value < 0.05, ** p-value < 0.01, *** p-value < 0.001.

### MSI1 promotes translation of recombinant *Gnat1* in NIH 3T3 cells

To demonstrate a direct activation of protein expression by Musashi we examined the effect of MSI1 on protein expression from *Gnat1* clones carrying full length 3’-UTR in a heterologous system, NIH 3T3 cells. The NIH 3T3 cells were chosen for the low levels of endogenous MSI1 and MSI2 protein expression. We created *Gnat1* clones that contained either wild type 3’-UTR or a mutant 3’-UTR in which the TAG sites were changed to TGA to prevent Musashi binding. The wild type and mutant clones carried HA and T7 epitope tags, respectively. The clones were mixed together in equal amounts and co-transfected in NIH 3T3 cells with an empty vector or vectors expressing GFP, *Pcbp2*, or *Msi1*. *Pcbp2* is used as negative control along with the empty vector and GFP. PCBP2 is a RNA binding protein that like MSI1 and MSI2 is abundantly expressed in photoreceptors, shuttles between the nucleus and the cytoplasm, and regulates splicing and translation (Ling et al., 2020; Makeyev and Liebhaber, 2002). The products of the wild type and mutant clones were distinguished by the attached HA (wild type) and T7 (mutant) epitope tags. The epitope tags were detected by antibodies on Western blot or by hydrolysable probes in multiplexed RT-qPCR, to measure protein and mRNA expression respectively. The effect of the co-transfected constructus on *Gnat1* expression was measured as the change in the ratio of the HA to the T7 signal.

One-way ANOVA found a significant effect of the co-transfected expression vector on the HA/T7 GNAT1 protein ratio (F(3,16)=20.52, p-value<0.0001). Tukey HSD post-hoc test showed that MSI1 has a significant (p-value<0.001) effect on the GNAT1 protein expression compared to the empty vector, or the vectors expressing GFP or PCBP2 (Figure 8C and D). The analysis of the *Gnat1* mRNA levels also revealed a significant effect of the co-transfected expression construct (one-way ANOVA F(3,20)=5.7, p-value=0.0055). The observed effect was due to a marginal increase in *Gnat1* transcript levels in response to MSI1 compared to empty vector and vector expressing PCBP2 (Figure 8D). The two fold increase in GNAT1 protein expression without a corresponding increase in transcript levels recapitulates the regulation of GNAT1 by Musashi that we observed in photoreceptor cells and points to a role for the Musashi proteins as activators of translation.

## Discussion

### MSI1 and MSI2 are essential and redundant in mature photoreceptor cells

Musashi was initially described as a protein in *Drosophila* that represses the translation of the *Notch* regulator *Numb* to maintain stem cell renewal a function that is preserved in vertebrates (Fox et al., 2015; Imai et al., 2001; Kharas et al., 2010; Li et al., 2015; Sakakibara et al., 2002). Expression of Musashi proteins is not confined to stem cells. MSI1 and MSI2 are expressed in various cell types, including mature neurons. We recently showed that MSI1 and MSI2 protein expression is exceptionally high in the retina and in photoreceptor cells (Sundar et al., 2020). Combined knockout of *Msi1* and *Msi2* in retinal and photoreceptor progenitor cells abolished the morphogenesis and function of the photoreceptor cells without significantly affecting the commitment of progenitors to the photoreceptor lineage (Sundar et al., 2020). Mice with single deletions of *Msi1* and *Msi2* in the developing retina exhibited an early reduction in vision and delayed retinal degeneration, respectively. The phenotype of retinal knockouts of *Msi1* and *Msi2* and an apparent switch from MSI1 to MSI2 protein expression during the period when photoreceptors elaborate their light sensing outer segment suggested separate roles for the two proteins in developing and mature photoreceptor cells.

Here we used an inducible knockout model to determine the function of the Musashi protein in mature photoreceptor cells. The phenotype of the combined *Msi1* and *Msi2* knockout mice demonstrated that the two proteins are critical for photoreceptor function and survival. Disruption of the two genes resulted in rapid loss of vision and retinal degeneration (Figure 2 and Figure 2 - figure supplement 1). The Musashi proteins were fully redundant in mature photoreceptors and the single *Msi1* or *Msi2* knockouts did not have detectable phenotypes. This is in contrast to our previous work where using *Six3-Cre* and *Nrl-Cre* models to delete the genes during retinal and rod photoreceptor development resulted in partial loss of vision and retinal degeneration (Sundar et al., 2020). It is unclear why *Msi1* and *Msi2* were only partially redundant during retinal development while they were fully redundant in mature photoreceptor cells.

While hundreds of mutations in dozens of genetic loci lead to loss of vision, to date no vision defects have been associated with mutations in the *Msi1* and *Msi2* genes despite their critical function in photoreceptor cells. The lack of vision defects associated with the Musashi genes is likely a combination of their redundancy and their critical role in stem cell maintenance. The redundancy of the two proteins ensures that mutations in one of the genes will be complemented by the other. At the same time, loss of both *Msi1* and *Msi2* will result in embryonic lethality and preempt the observation of retinal phenotypes.

### MSI1 and MSI2 activate inclusion of alternative exons *in vivo*

We previously showed that photoreceptor cells express a distinct splicing program that utilizes a large number of microexons. Motif enrichment analysis suggested that the inclusion of photoreceptor-specific exons is driven by the Musashi proteins (Murphy et al., 2016). Recent studies on mouse tissues, flow sorted retinal cells, and human retinal organoids showed a similar photoreceptor-specific splicing program directed by Musashi (Ling et al., 2020; Ottaviani et al., 2021). Here we demonstrate that the Musashi proteins directly regulate splicing *in vivo* in the retina. Binding of the Musashi proteins downstream of photoreceptor specific alternative exons promotes their inclusion in the mature transcripts. The Musashi proteins promote the inclusion of more than half of the exons we previously defined as photoreceptor specific (Murphy et al., 2016). This leaves a sizable population of photoreceptor-specific exons that rely on other factors for their splicing. Recent studies have highlighted two such factors that either cooperate with Musashi (PCBP2) or promote the splicing of independent sets of exons (SRRM3) (Ciampi et al., 2021; Ling et al., 2020).

In addition to the exons activated by the Musashi proteins, a comparable number of exons appeared to be repressed by them. The repressed exons do not show a significant pattern of enrichment of MSI1 binding to them or to the adjacent introns. It is possible that the Musashi proteins may have more than one mode of directly repressing splicing and our dataset does not have sufficient power to detect these interactions. It is also likely that many of the repressed exons are not direct targets of the Musashi proteins, but are regulated by factors whose expression is controlled by MSI1 and MSI2. For example, the Musashi proteins negatively regulate the expression of the SFRS9 protein (Supplementary Table 4). Consequently, the splicing of exons dependent on SRSF9 will be repressed indirectly by the expression of the Musashi proteins in photoreceptors.

### Dispensable conserved microexons

Use of alternative microexons is a hallmark of the nervous system. The majority of these alternative exons are conserved which has led to the obvious hypothesis that they perform essential functions in neurons. For several microexons such function has been demonstrated *in vivo* in animal knockout models (Gonatopoulos-Pournatzis et al., 2020; Johnson et al., 2019; Lin et al., 2020; Nakano et al., 2018). We reasoned that exons that are conserved in vertebrates and are specifically spliced in photoreceptor cells will likely have an important role in vision. We deleted four such conserved exons in the *Ttc8*, *Cep290*, *Cc2d2a* and *Cacna2d4* genes, and one exon in the *Slc17a7* gene that is present only in rodents and is potentially nonessential (Figure 6 - figure supplement 1). All five exons are photoreceptor-specific and are included in nearly all of the transcripts from the corresponding genes in photoreceptor cells (Figure 6A) (Ling et al., 2020; Murphy et al., 2016). The genes hosting four of the exons, *Ttc8*, *Cep290*, *Cc2d2a* and *Cacna2d4*, are essential for vision (Bachmann-Gagescu et al., 2012; Dilan et al., 2018; Rachel et al., 2012; Wycisk et al., 2006). To our surprise the exon knockout animals did not show a detectable phenotype. A triple knockout of the exons in *Ttc8*, *Cep290*, and *Cc2d2a*, all essential components of the cilium, also failed to manifest an adverse phenotype. The lack of phenotype in our exon knockout animals raises questions about the nature of the selective pressures that have led to the conservation of these exons. It is possible that the selective pressures are exerted by factors that are absent from the environment under which laboratory mice are reared. An alternative explanation is that these exons do not have function and do not alter the properties of the proteins. The conservation of such functionally neutral exons will be due to elimination of mutations that can negatively affect the function of the host protein. Functionally neutral exons will be under purifying selection that tolerates wholesale deletion of the exon. In support of this model, the photoreceptor-specific exons in *Ttc8* and *Cc2d2a* are absent from certain species (Figure 6 - figure supplement 1). While determining the reasons for the sequence conservation of the microexons in the *Ttc8*, *Cep290*, *Cc2d2a*, and *Cacna2d4* genes will require further work, our results caution against using sequence conservation as a sole indicator of function.

### MSI1 and MSI2 promote protein expression in photoreceptor cells

Musashi proteins can both activate and repress protein translation. In flies and cultured mammalian cell lines the Musashi proteins repress translation of *Numb* and p21^Waf1/Cip1^, while in frog oocytes they activate early translation of Mos and Cyclin B5 after progesterone stimulation (Arumugam et al., 2010; Battelli et al., 2006; Charlesworth et al., 2006; Imai et al., 2001). Furthermore, recent genome wide studies integrating Ribo-Seq and CLIP-Seq data showed that, within the same cell, MSI1 and MSI2 repress translation of certain transcripts while activating others (Karmakar et al., 2021; Katz et al., 2014). The transcriptome-wide studies also show that translation of relatively few of the large numbers of transcripts that are bound by the Musashi proteins is affected when the levels of MSI1 and MSI2 are manipulated (Karmakar et al., 2021).

Here we present an integrative analysis of the effect of MSI1 and MSI2 on protein translation in photoreceptor cells *in vivo*. A limitation of our work is that while we knocked out *Msi1* and *Msi2* specifically in photoreceptor cells, the proteomics, RNA-Seq, and eCLIP-Seq experiments were performed on the entire retina. To isolate the signal derived from photoreceptor cells, we focused our analysis on transcripts highly expressed in photoreceptors compared to other retinal cells and relied on the fact that photoreceptor cells are the dominant cell type in the retina comprising approximately half of the cells in that tissue. As we are excluding from our analysis transcripts that are expressed in other cells of the retina at levels comparable to those in photoreceptors, we derived a broad but far from comprehensive picture of the effect of MSI1 and MSI2 on the transcriptome and proteome of photoreceptor cells.

Regardless of this limitation, we demonstrate that the combined deletion of *Msi1* and *Msi2* alters protein expression from a set of transcripts without significantly affecting the levels of these transcripts. In agreement with the previously published transcriptome-wide studies, our integrative analysis of eCLIP-Seq, RNA-Seq and proteomics data show that the levels of relatively few proteins are affected in the Musashi knockout mice, compared to the thousands of transcripts bound by MSI1 at their 3’-UTRs.

In photoreceptors, the Musashi proteins act largely to promote protein expression. There are only two exceptions, PROM1 and IMPG2, where the combined knockout of *Msi1* and *Msi2* resulted in elevated protein levels. This is an unexpected finding in light of the canonical view of the Musashi proteins as repressors of translation. While the Musashi proteins are known to promote translation, our study is the first to show a broad role for the Musashi proteins as almost exclusive post-transcriptional activators of protein expression. Previous studies in neuronal precursors and K562 cells have shown a more balanced effect with comparable numbers of proteins being activated and repressed by MSI1 and MSI2 (Karmakar et al., 2021; Katz et al., 2014). Further studies will be required to determine the molecular mechanism by which MSI1 and MSI2 promote protein expression in photoreceptor cells. Nevertheless, we show that the translation of at least one of the targets identified in this work, *Gnat1*, can be stimulated by MSI1 in a heterologous system.

The proteins regulated by MSI1 and MSI2 are central to the function and survival of photoreceptor cells such as components of the outer segment (GNAT1, CNGA1, PRCD, ROM1), and chaperones responsible for the folding of proteins critical for photoreceptors (AIPL1, PDC). Photoreceptor cells need to produce high levels of these proteins in order to replace the photoreceptor outer segments every 10 days. This renewal process does not reuse proteins already present in the outer segment. Instead, new membranes and proteins are delivered to the bottom of the outer segment stack, while old segments are phagocytosed and digested by retinal pigmented epithelium from the top of the stack. Reduced rate of production of outer segment proteins and the chaperones that fold them will impede the outer segment renewal process leading to loss of vision and degeneration of the photoreceptor cells.

## MATERIALS AND METHODS

### Animals

All animal experiments were conducted with the approval of the Institutional Animal Care and Use Committee at West Virginia University. Both males and females were used in all experiments. The mouse lines in this study were in the C57BL6/J background and devoid of the naturally occurring *rd1* and *rd8* alleles. The mice were genotyped at weaning unless otherwise specified in the results section. The primers used for genotyping of the targeted alleles are listed in Supplementary Table 14.

Mice carrying *Msi1 ^flox/flox^* and *Msi2 ^flox/flox^* were provided by Dr. Christopher Lengner from the University of Pennsylvania (Li et al., 2015; Park et al., 2014). Mice carrying the floxed alleles were crossed with *Pde6g-Cre^ERT2^* mice to enable photoreceptor-specific conditional knockout of *Msi1* and *Msi2* (Koch et al., 2015). The conditional deletion of *Msi1* and *Msi2* in mature photoreceptor cells was induced by intraperitoneal injection of tamoxifen in corn oil (Sigma-Aldrich catalog #T5648-1G) at concentrations of 100 mg/kg body weight for three consecutive days.

The knockouts of the photoreceptor specific exons in *Ttc8*, *Cep 290*, and *Cc2d2a* were created using CRISPR/Cas9. Two guide RNAs targeted at sites upstream and downstream of each alternative exon were used to cause full deletion of the exon and the proximal parts of the introns. The guide RNAs were synthesized by Synthego and IDT. The guide RNA targeting sequences are listed in Supplementary Table 13. The guide RNAs and Cas9 (Thermo Fisher) were assembled into ribonucleoprotein complexes and electroporated into zygotes by the WVU transgenic core facility. The founders were back-crossed to C57BL6/J mice (Jackson Laboratory) for 5 generations. To map the borders of the deletions, the exon knockout alleles were amplified by PCR using the genotyping primers and sequenced by Sanger sequencing (Figure 6 - figure supplement 2).

Adult zebrafish (*Danio rerio*) animals were maintained at 28°C with standard 14/10 light/dark cycles. For dissections, we used adult Tubingen long-fin strain (approximately 22 months old). Equal female and male zebrafish were euthanized in an ice bath of system water until the termination of buccal and gill motion. Tissue dissection was performed in physiological saline E3h media (5 mM NaCl, 0.17 mM KCl, 0.33 mM CaCl2, 0.33 mM MgSO4, and 1 mM HEPES, pH 7.3). Collected samples were immediately collected in centrifuge tubes and frozen.

### Clones, cell lines, and transfection

A full length *Gnat1* clone (accession BC058810) was obtained from Horizon Discovery and recloned in pcDNA3.1(+) (Invitrogen). A matching clone in which all TAG triplets in the 3’-UTR were mutated to TGA to disrupt the Musashi binding sites was created using gene synthesis (Genscript). Gibson assembly was used to reclone the cDNAs into pcDNA3.1(+) vector and attach HA- and T7-tags to the wild type and mutant clone, respectively. Full length *Msi1* clone with N-terminal Flag epitope tag in pcDNA3.1 was described before (Murphy et al., 2016). Full length, codon optimized mouse Pcbp2 clone with N-terminal Flag epitope tag was produced by gene synthesis (Genscript) and cloned in pcDNA3.1 (Invitrogen).

NIH 3T3 cells were grown in DMEM supplemented with 10% Fetal Bovine Serum. The cells were maintained in a humidified incubator at 37°C in 5% CO_2_ atmosphere. Transfection with polyethyleneimine was carried out as described before (Boussif et al., 1995). Briefly, the 6 hours prior to transfection the cells were seeded at 3.2*10^5^ cells per well in 6 well plates. A total of 1ug of DNA was used per transfection, containing 250 ng of each wild type and mutant *Gnat1* construct, and 500ng of expression vector that was either empty or carried a GFP (pAcGFP N1, Clontech), *Pcbp2* or *Msi1* clone. The cells were collected to analyze protein and mRNA expression.

### RNA extraction and RT-PCR

RNA was extracted with Trisol and precipitated with isopropanol. The RNA was then dissolved and treated with RNAse-free DNase I (Roche) for 20 minutes at 37°C. After DNA digestion the reactions were extracted once with chloroform and the RNA was precipitated with ethanol. RT-PCR analysis of alternative splicing using fluorescently labeled primers was described before (Murphy et al., 2016; Percifield et al., 2014).

The levels of Gnat1 transcripts expressed from the recombinant clones in 3T3 cells were determined by multiplexed RT-qPCR. Hydrolysis probes to the HA and T7 tags were used to detect Gnat1 transcripts with wild type and mutant 3’-UTRs, respectively. The RT-qPCR was performed using Luna One Step RT-qPCR mix with dUDG (NEB). Amplification using Luna One Step qPCR mix with UDG (NEB) that did not contain a reverse transcriptase component (NEB) served as no-reverse -transcriptase controls for DNA contamination. The ratio of the transcript levels measured by the HA and T7 probes was used to determine the effect of each treatment on the mRNA levels expressed from the constructs carrying wild type and mutant 3’-UTRs. The primers and probes used for alternative splicing and RT-qPCR analysis are listed in Supplementary Table 14.

### Electroretinography (ERG) Measurement and Preparation of Animal

ERGs were measured using either UTAS Visual Diagnostic System with Big-Shot Ganzfeld device (LKC Technologies, Gaithersburg, MD, USA) or Celeris system with Espion software (Diagnosis LLC, Lowell, MA, USA). Prior to testing, mice were dark-adapted overnight. All further handling of mice following dark adaptation was performed under deep red illumination. The mice were anesthetized by inhalation of 1.5% isoflurane mixed with 100% oxygen at a flow rate of 2.5 l/min. The pupils were topically dilated with a drop of tropicamide and phenylephrine-hydrochloride, allowing drops to sit on both eyes for 10 mins. After that, mice were transferred to a heated platform connected to a nose cone that allows for a continuous flow of isoflurane. A reference electrode was inserted sub-dermally between the eyes of the mouse, and ERG responses were collected from both eyes using wire electrodes placed on the center of each cornea, with contact being made using a drop of 0.3% Hypromellose solution. To deliver the stimulus, a Ganzfeld Bowel was used with LED white arrays at increasing intensities. Dark-adapted scotopic photoresponse was recorded under the dim red light using a single LED white light flashes of luminescence ranging from 2.45**·**10^-4^ to 2.4 cd-s/m^2^. For photopic response, animals were light-adapted for 10 min in the presence of rod-saturating 30 cd-s/m^2^ ambient white light prior to recording the photopic response.

### Western Blot

Mouse retinas were lysed using RIPA buffer (50 mM Tris HCl-pH 8.0, 150 mM NaCl, 1.0% TritonX-100, 0.5% sodium deoxycholate, 0.1% sodium dodecyl sulfate) supplied with protease (Sigma-Aldrich catalog# 535140-1ML) and phosphatase inhibitors cocktail (Sigma-Aldrich catalog # P5726-1 ML). After homogenization, the lysate was incubated on ice for 10 mins, then cleared by centrifugation for 15 mins. 20 μg of protein extract was resolved in 4-20% polyacrylamide SDS–PAGE gel and transferred onto polyvinylidene difluoride (PVDF) membranes (Immunobilon-FL, Millipore). After blocking with BSA in PBST (Phosphate-buffered saline with 0.1% Tween-20), the membranes were blocked and probed with primary antibodies overnight at 4 °C, followed by incubation with fluorescently labeled (Alexa Fluor 647 or 488, Jackson ImmnuoResearch) secondary antibodies for 1 hour at room temperature. The membranes were then scanned on Amersham Typhoon Phosphorimager (GE Healthcare). GFP protein in transfection samples was detected by its intrinsic fluorescence in the 488nm channel. To quantify the protein expression across membranes the band intensities detected by each antibody were scaled to the median signal for the membrane. The scaled expression values were then normalized to the scaled values of the corresponding controls (loading controls, or in the transfection experiments to proteins expressed from co-transfected constructs). The antibodies used for western blot analysis are listed in Supplementary Table 12.

### Retinal histology

The whole eyecups from the knockout and control mice were enucleated. The eyes were then fixed using a Z-fixative (Excalibur Pathology Inc). Tissue processing, including paraffin embedding and hematoxylin and eosin (H&E) staining was performed at Excalibur Pathology. Images of the stained slides were collected using a Nikon Brightfield Microscope operated by Element software (Nikon). To evaluate the photoreceptor cell loss, we counted the number of nuclei within the outer nuclear layer (ONL) using the NIS elements software. The counting was done at ten equidistant locations centered on the optical nerve and moving toward the periphery in 400 µm increments. Five locations were on the inferior side (-5 to -1) and five on the superior side (1 to 5) of the retina relative to the optic nerve. For each location and the number of nuclei reported is the average of 4 technical replicates. The nuclei counts were averaged over 3 biological replicates that represent retinas from three separate animals.

### Immunocytochemistry

Eyes were enucleated, and small window was cut in the cornea before immersing it in 4% paraformaldehyde fixative (4% PFA in PBS: 137 mM NaCl, 2.7 mM KCl, 10 mM Na_2_HPO_4_, and 1.8 mM KH_2_PO_4_, pH 7.2) for 3 hours on a rotator. Eyecups were dehydrated by sequential incubation in 7.5%, 15%, and 22% sucrose in 1xPBS. Eyecups were then snap-frozen in optimal cutting temperature compound (OCT) blocks. Serial 16 µm sections were cut on a Leica CM1850 cryostat and mounted onto Superfrost Plus microscope slides (Fischer Scientific). Mounted retinal sections were washed 3 times for 10 mins each with PBS and then blocked with PBST for 1 hour (10% goat serum, 0.3% Triton X-100, and 0.02% sodium azide in PBS). Retinal sections were incubated overnight at 4°C with primary antibodies diluted in PBST supplemented with 5% goat serum. After three 15 min washes with PBST the sections were incubated for one hour with secondary antibodies diluted 1:1000 in PBST supplemented with 5% goat serum and 4′,6-diamidino-2-phenylindole. The sections were washed three times for 15 min with PBST, mounted with Prolong Antifade reagent (Thermofisher) and secured with coverslips. The sections were imaged on a Nikon C2 laser scanning confocal microscope. The laser power, gain and offset settings were maintained the same when imaging sections from knockout and control littermates. The antibodies used for immunofluorescence staining are listed in Supplementary Table 12.

### RNA sequencing

Total RNA was isolated at day 21 post tamoxifen injection using Tri-reagent (Sigma) from retinas in four biological replicates of *Msi1^+/+^/Msi2^+/+^* and *Msi1^-/-^/Msi2^-/-^* mice. Sequencing libraries were prepared by the West Virginia University genomics core using KAPA Hyper RNA with Riboerase (Roche). The libraries were sequenced by the University of Illinois DNA services core on Illumina HiSeq 4000 at average depth of 44 million 100nt paired end reads. The RNA-Seq data is available at the NCBI Sequence Read Archive under project accession PRJNA795137.

RNA-Seq reads were aligned to the mouse genome (GRCm38) using HISAT2 (Kim et al., 2019). The mapped reads were summarized using Rsubread and differential gene expression analysis was carried out by edgeR (Supplementary Table 8) (Liao et al., 2019; Robinson et al., 2010). Inclusion levels of cassette exons were calculated by rMATS (4.1.0), using reads spanning exon-exon junctions (Shen et al., 2014).

### CLIP-sequencing and meta exon analysis

The rabbit anti-MSI1 (1:1000; catalog# ab 52865, Abcam, Cambridge, MA) was used for eCLIP. Briefly, retinas from wild-type mice (80 mg per replicate) were collected and placed in ice-cold PBS in a 10 cm^2^ plate. Plates containing retinas were then placed on ice and UV-crosslinked (254 nm, 200 mJ/cm^2^ ) using UV Stratalinker^TM^ 2400. UV-crosslinked retinas were then snap frozen in liquid nitrogen. Further tissue processing, eCLIP library prep, and sequencing were carried out by Eclipse Bioinnovations following a previously published protocol (Van Nostrand et al., 2016). The raw data obtained from the eCLIP-Seq are available at the NCBI Sequence Read Archive under project accession PRJNA795195.

To analyze the raw eCLIP data, the adapter sequences were first trimmed using cutadapt (Martin, 2011). HISAT2 was used to map the reads to version GRCm38 of the mouse genome and the mapped reads were deduplicated by umi-tools using the unique molecular identifier (UMI) barcodes built into the adapters (Kim et al., 2019; Smith et al., 2017). Crosslink sites were identified and clustered (Supplemental data set 1) into regions using PureClip (Krakau et al., 2017). Motifs enriched in the 51 nucleotide sequence fragments centered on the crosslink sites were identified by HOMER and DREME (Bailey, 2011; Heinz et al., 2010).

Meta-exon analysis was performed using the RBP-maps software package on non-redundant sets of alternatively spliced exons identified in our RNA-Seq analysis of *Msi1*/*Msi2* double knockout in photoreceptor cells (Yee et al., 2019). The distribution of MSI1 crosslinks around exons downregulated or upregulated in the photoreceptor specific *Msi1*/*Msi2* double knockout was compared to alternative exons that were not affected by the knockout (Supplementary Tables 9, 10, and 11). 1000 random permutations of the non-regulated exon set were used to determine the 99.5% confidence intervals as described by Yee at al (Yee et al., 2019).

### Proteomics analysis

Retina samples from *Msi1^+/+^/Msi2^+/+^* and *Msi1^-/-^/Msi2^-/-^* were collected 21 days after tamoxifen injections. Five biological replicates were used for each wild-type and knockout group. Tissue processing and proteomics quantification of snapped frozen retina samples was performed by IDeA proteomics. Briefly, proteins were reduced, alkylated, and purified by chloroform/methanol extraction prior to digestion with sequencing grade modified porcine trypsin (Promega). Tryptic peptides were labeled using tandem mass tag isobaric labeling reagents (Thermo) following the manufacturer’s instructions and combined into one 10-plex sample group. The labeled peptide multiplex was separated into 46 fractions on a 100 x 1.0 mm Acquity BEH C18 column (Waters) using an UltiMate 3000 UHPLC system (Thermo) with a 50 min gradient from 99:1 to 60:40 buffer A:B ratio under basic pH conditions, and then consolidated into 18 super-fractions. Each super-fraction was then further separated by reverse phase XSelect CSH C18 2.5 um resin (Waters) on an in-line 150 x 0.075 mm column using an UltiMate 3000 RSLCnano system (Thermo). Peptides were eluted using a 60 min gradient from 98:2 to 60:40 buffer A:B ratio. Eluted peptides were ionized by electrospray (2.2 kV) followed by mass spectrometric analysis on an Orbitrap Eclipse Tribrid mass spectrometer (Thermo) using multi-notch MS3 parameters. MS data were acquired using the FTMS analyzer in top-speed profile mode at a resolution of 120,000 over a range of 375 to 1500 m/z. Following CID activation with normalized collision energy of 35.0, MS/MS data were acquired using the ion trap analyzer in centroid mode and normal mass range. Using synchronous precursor selection, up to 10 MS/MS precursors were selected for HCD activation with normalized collision energy of 65.0, followed by acquisition of MS3 reporter ion data using the FTMS analyzer in profile mode at a resolution of 50,000 over a range of 100-500 m/z. Buffer A is 0.1% formic acid, 0.5% acetonitrile; Buffer B is 0.1% formic acid, 99.9% acetonitrile. Both buffers were adjusted to pH 10 with ammonium hydroxide for offline separation.

To create a database of proteins expressed in the retina we first filtered our mouse retina RNA-Seq data to remove genes with median expression across all samples that were below the median expression for the dataset. As a result we selected 15,626 genes with expression equal or more than 1.2 RPKM. Ensembl release 79 was queried for annotated proteins produced by these genes resulting in a database of 34,055 protein sequences. Peptide identification against the retinal protein database was performed using MS-GF+ (version v2021.03.22) with parent ion tolerance of 10 ppm, a reporter ion tolerance of -0.00335 Da and +0.0067 Da, and requiring fully tryptic peptides (Kim and Pevzner, 2014). Only peptides with peptide level Q-value of 0.05 or below were accepted. The MSnbase package from R/Bioconductor was used to quantify the MS3 reporter ions and combine the identification and quantification data (Gatto et al., 2021; Gatto and Lilley, 2012). Differential protein expression analysis was performed using the DeqMS package from R/Bionconductor (Zhu et al., 2020). Protein changes with adjusted p-value below 0.05 and fold change of more than 1.5 were considered significant.

WebGestalt (WEB-based Gene SeT AnaLysis Toolkit) was used to perform enrichment analysis on the Gene Ontology and KEGG databases for the proteins with significant changes in gene expression. Only terms enriched at FDR<0.05 were reported (Wang et al., 2013). The mass spectrometry proteomics data have been deposited to the ProteomeXchange Consortium via the PRIDE partner repository with the dataset identifier PXD030748 and 10.6019/PXD030748 (Deutsch et al., 2020; Perez-Riverol et al., 2019).

### Experimental design and Statistical analysis

Age-matched males and females in the C57BL6/J background were used in all experiments. The statistical analysis and data visualization was done using GraphPad Prism and R/Bioconductor. Unpaired Student’s t-test was used to assess statistical significance between control and knockout samples. Statistical significance was determined with one-way or two-way ANOVA followed by pairwise comparisons as indicated in the text. All data were presented as the mean ±standard error of the mean (SEM).

## Supporting information

Supplemental Tables 1, 2, and 3. Alternative splicing analysis

Supplemental Table 4. Differential protein expression.

Supplemental Tables 5 and 6. Gene ontology and pathway enrichment analysis.

Supplemental Table 7. Proteomics and RNA-Seq data related to Figure 8

Supplemental Table 8. Differential gene expression.

Supplemental Tables 9, 10, and 11. Alternative splicing analysis

Supplemental Tables 12, 13, and 14. Antibodies, guide RNAs and primers.

Supplemental data set 1. Bed files (GRCm38) with MSI1 binding sites and regions identified by eCLIP

## ACKNOWLEDGEMENTS

We are grateful to Dr. Christopher Lengner for the generous donation of the *Msi1^fl/fl^* and *Msi2^fl/fl^* mice. We thank Drs. Roberta Leonardi and Aaron Robart for critical reading of this manuscript.

## FIGURES

**Figure 1 - Figure supplement 1.**
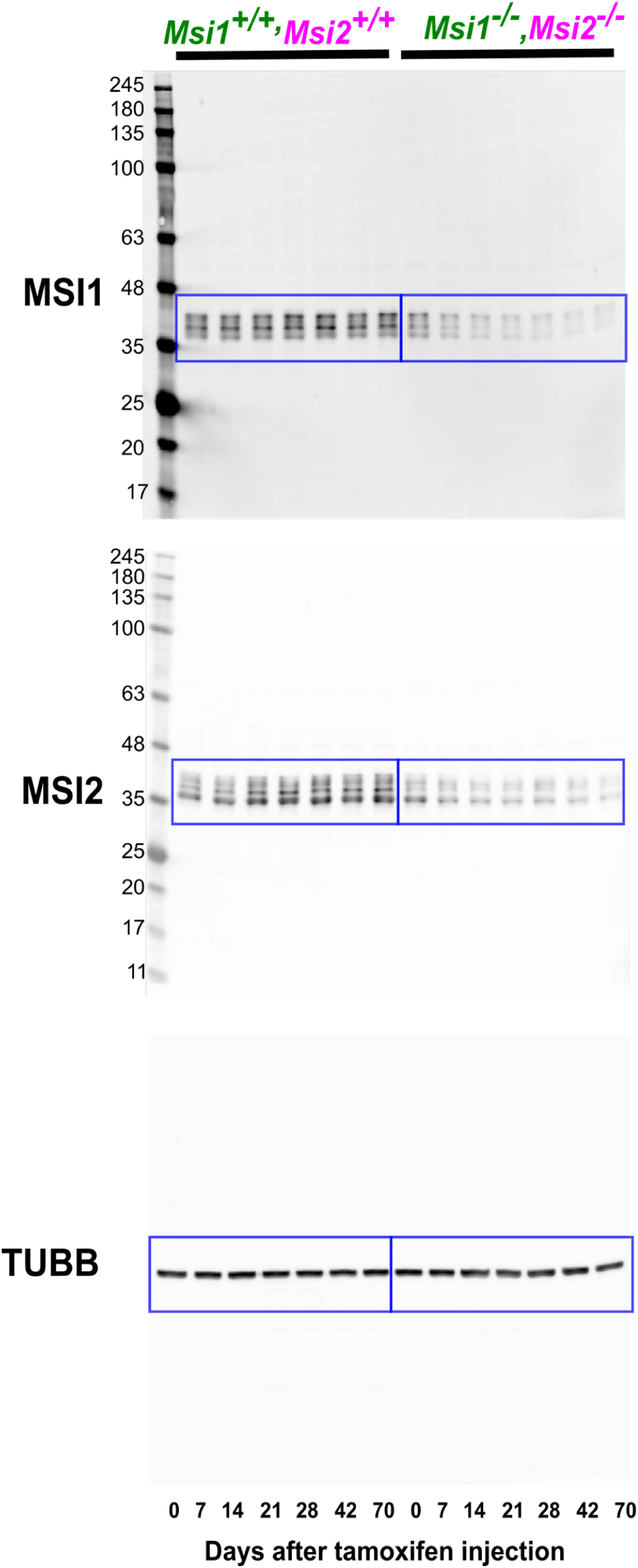
Full size western blot images for the data presented on Figure 1B (blue boxes).

**Figure 2 - figure supplement 1.**
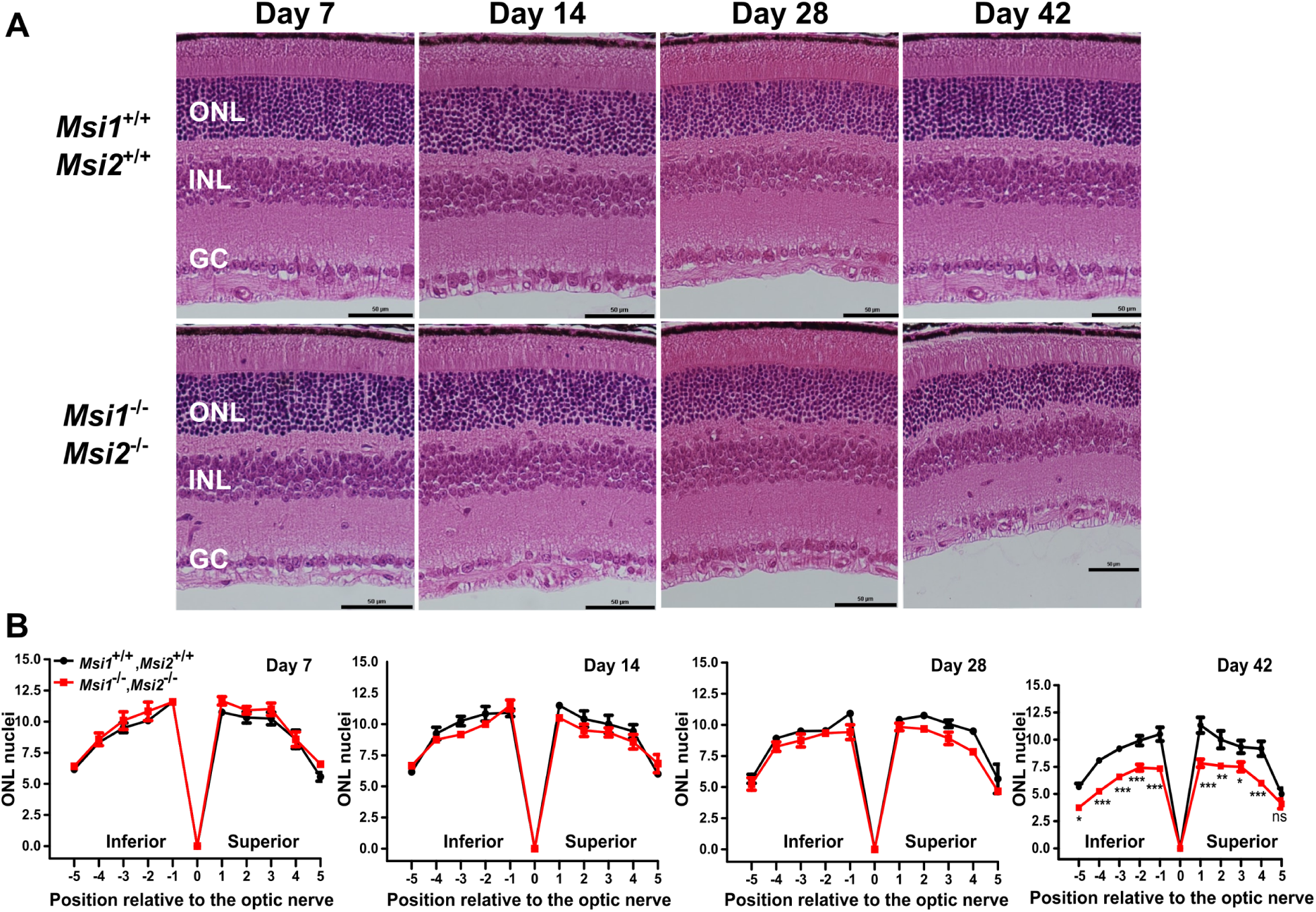
Outer nuclear layer thickness of the retina after double knockout of *Msi1* and *Msi2* in photoreceptor cells. **(A)** Representative H&E-stained eye sections from the double *Msi1*/*Msi2* knockout and age matched controls at 7, 14, 28 and 42 days after inducing the knockout. ONL: outer nuclear layer (Photoreceptor nuclei), INL: inner nuclear layer, GC: ganglion cells. 40X objectives and scale bar represents 50 μm. (**B**) Spider plots displaying the thickness of the ONL as the number of nuclei measured at ten points stepped by 400µm from the optical nerve at different time points post-tamoxifen injection. Data are shown as mean ± SEM. Pairwise t-test with Bonferroni correction for multiple comparisons was used to determine the effect of the genotype on the outer nuclear layer thickness at each time point. Significance levels of the pairwise comparisons is indicated as: * p-value < 0.05, ** p-value < 0.01, *** p-value < 0.001.

**Figure 3-figure supplement 1.**
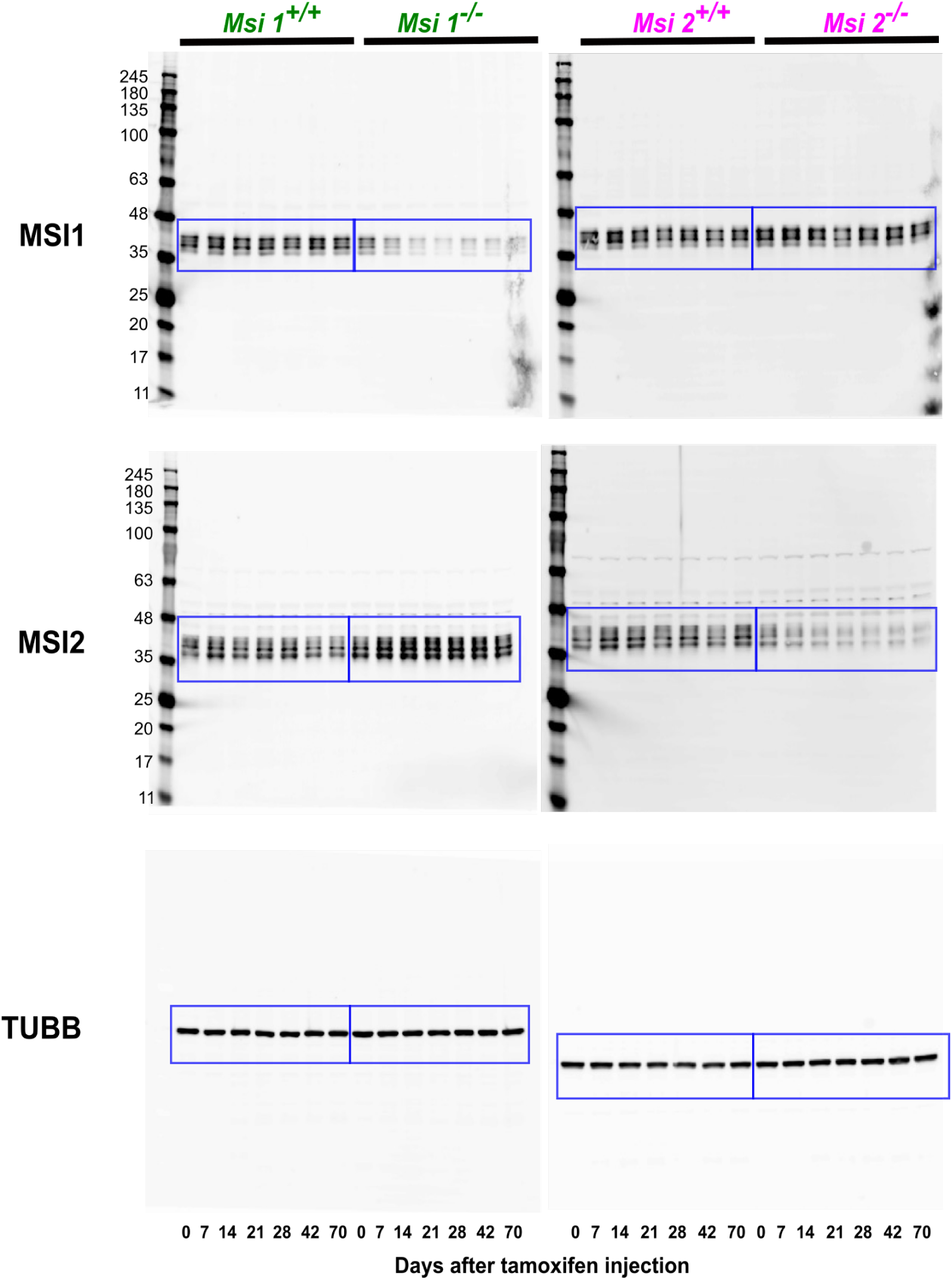
Full size western blot images for the data presented on Figure 1B (blue boxes).

**Figure 4 - figure supplement 1.**
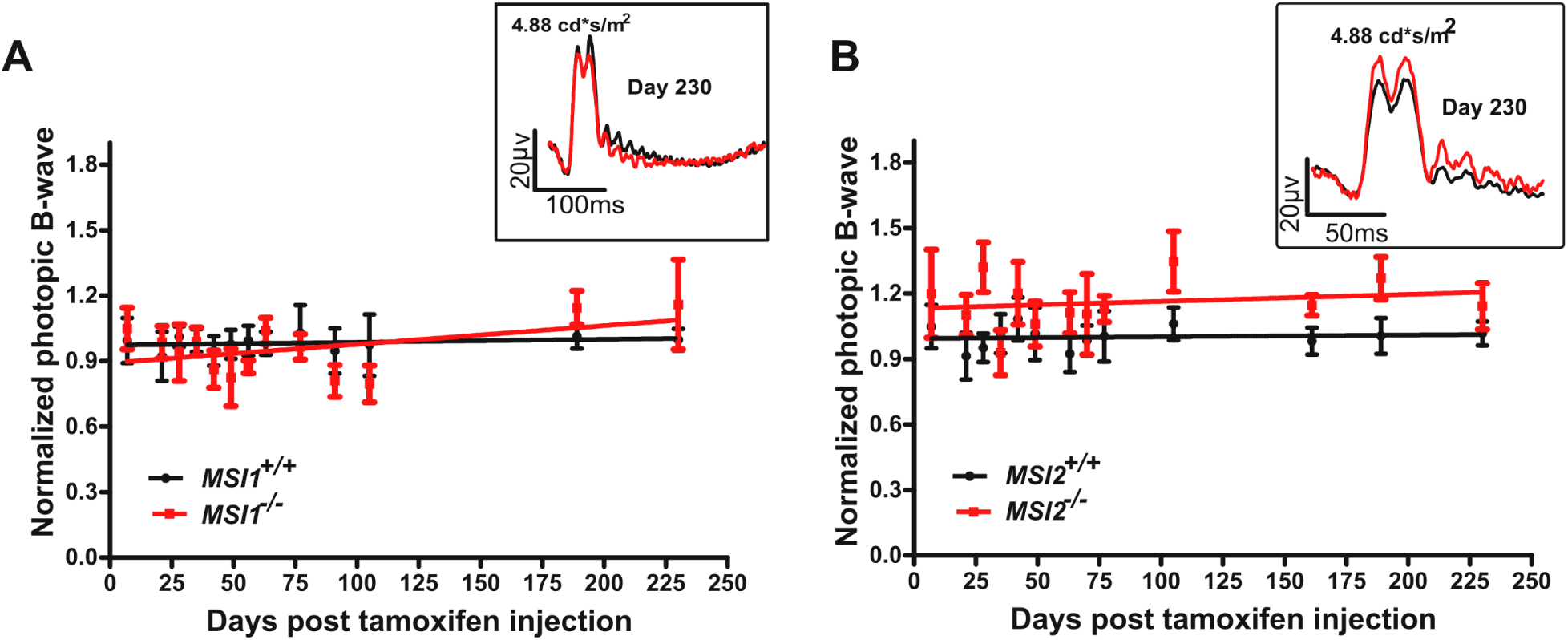
Normal photopic response to light in the single *Msi1* or *Msi2* knockouts. **A**) Photopic mean B-wave response of knockout (red) *Msi1^-/-^* **(A)** and *Msi1^-/-^* **(B)** mice (red line) and age matched controls (black) between day 16 and day 230 post tamoxifen injection. Photopic waveforms were obtained after light adaptation using 4.88 cd-s/m^2^ flashes. The insets show representative photopic (light-adapted) electroretinograms recorded 230 days post-injection using 4.88 cd-s/m^2^ flashes. The data points from the photopic responses of the single depletion of *Msi1* or *Msi2* are represented as mean ± SEM of 8 eyes, from 4 animals.

**Figure 5 - figure supplement 1.**
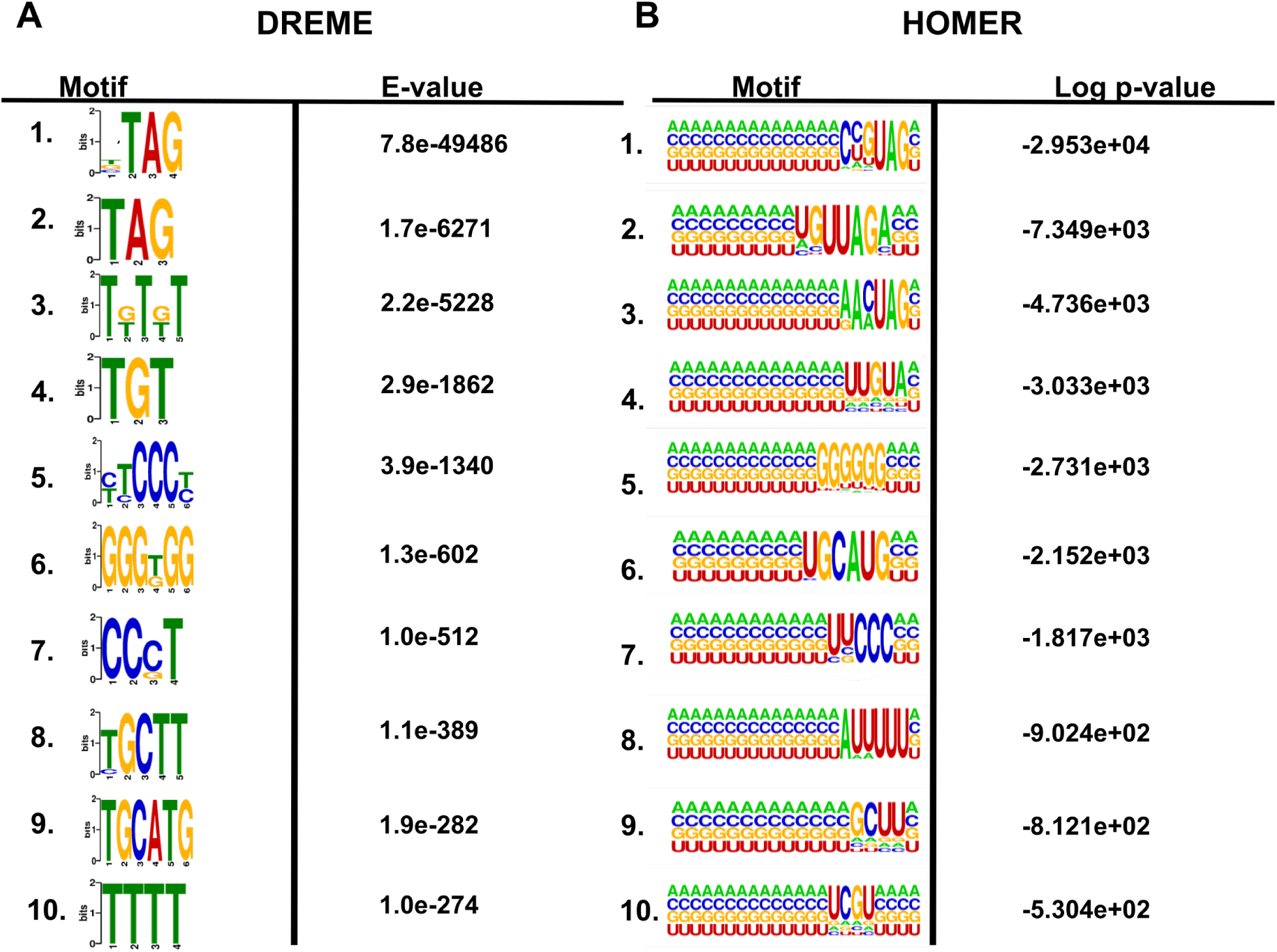
Sequence motifs enriched near eCLIP-Seq derived MSI1 crosslinks sites. Logos of the top ten significantly enriched motifs identified by DREME **(A)** or HOMER **(B)** in the vicinity of MSI1 crosslink sites.

**Figure 6 - figure supplement 1:**
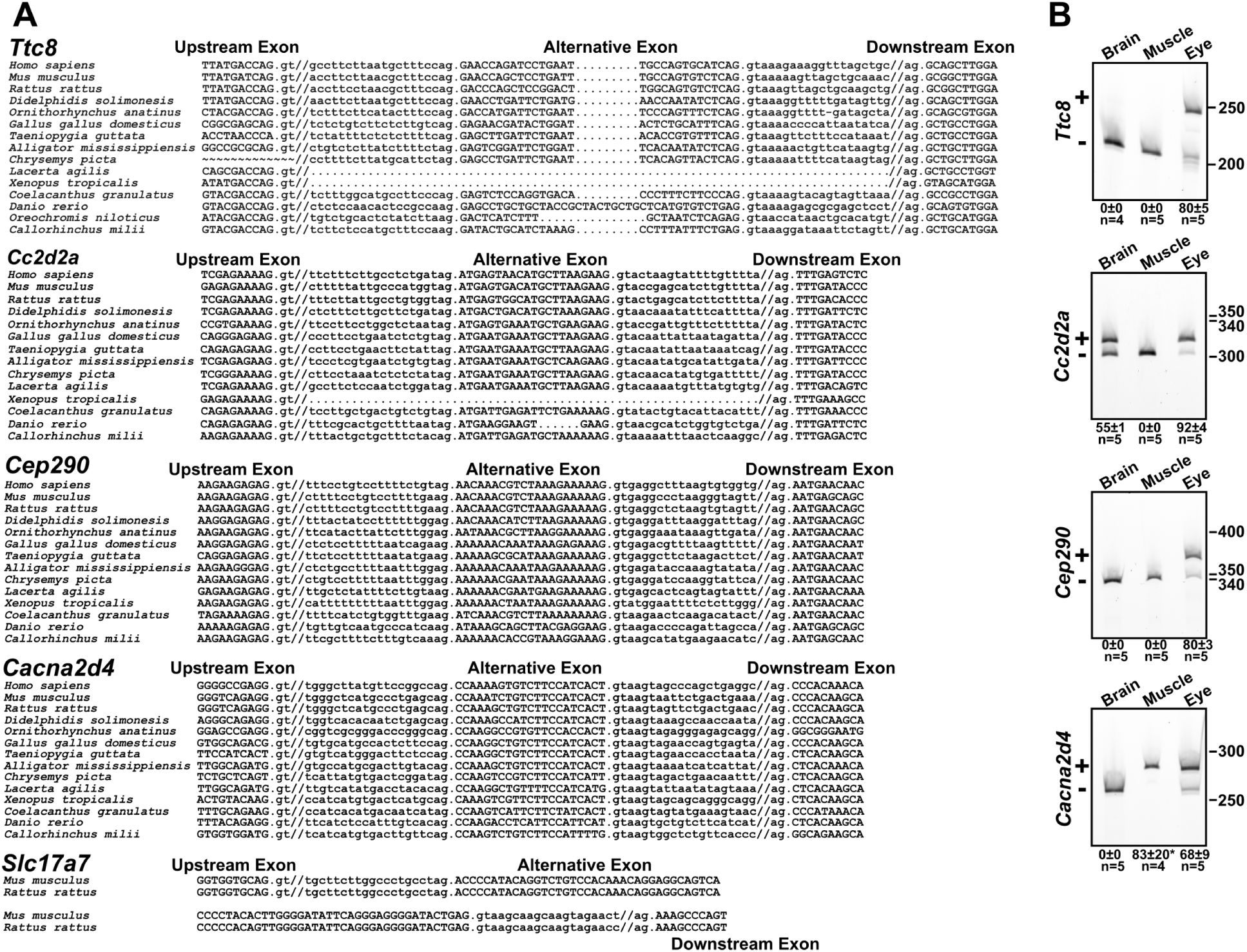
Conservation of the photoreceptor-specific exons of *Ttc8, Cc2d2a, Cep290, Cacna2d4, and Slc17a7* across vertebrates. **(A)** Alignments showing the alternative exons and parts of the flanking constitutive exons. Introns are in lower case and exons are in upper case. Forward slashes indicate where intronic sequences were removed from the alignment for ease of presentation. Homologous exons for the photoreceptor specific exons in *Ttc8, Cc2d2a, Cep290,* and *Cacna2d4* can be traced down to Chondrichthyes. The exons in *Ttc8,* and *Cc2d2a* can vary in length while preserving the reading frame or be completely absent from certain species. The upstream exon is not available for *Chrysemys picta* due to gaps in the genome sequence. The exon in *Slc17a7* is present only in rodents. **(B)** Analysis of the inclusion rate of the zebrafish homologues of the photoreceptor-specific exons in the *Ttc8*, *Cc2d2a*, *Cep290*, and *Cacna2d4* in brain, muscle, and eye samples. Numbers under the figure indicate the percent inclusion of the exon ± SEM. All four exons have high inclusion rates in the eye. Unlike their mouse homologues the zebrafish exons in the *Cc2d2a* and *Cacna2d4* genes are also included at high rate in the brain and muscle, respectively. *The exon in *Cacna2d4* was typically included at 100% (3 out of 4 tested samples).

**Figure 6 - figure supplement 2.**
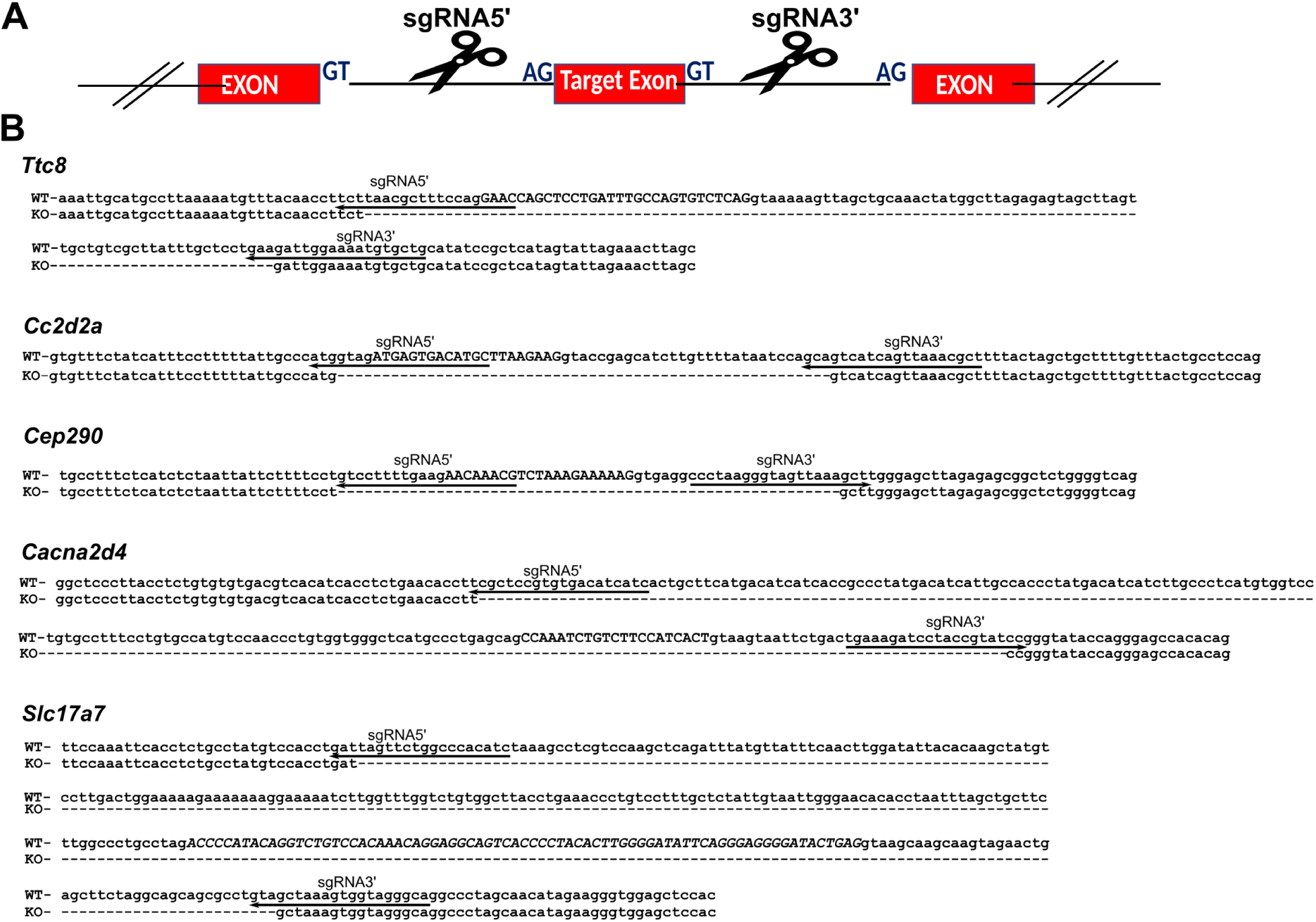
Exon deletion alleles generated by CRISPR/Cas9 mutagenesis. **A)** Schematic of the CRISPR/Cas9 targeting strategy. Two guide RNAs are used to direct cuts on both sides of the exon leading to its deletion. **(B)** Sequences of the knockout alleles (KO) aligned to the wild type genomic sequence (WT). The exons are shown in uppercase and introns are in lowercase. Arrows indicate the position and orientation (5’ to 3’) of the guide RNAs.

**Figure 7 - figure supplement 1.**
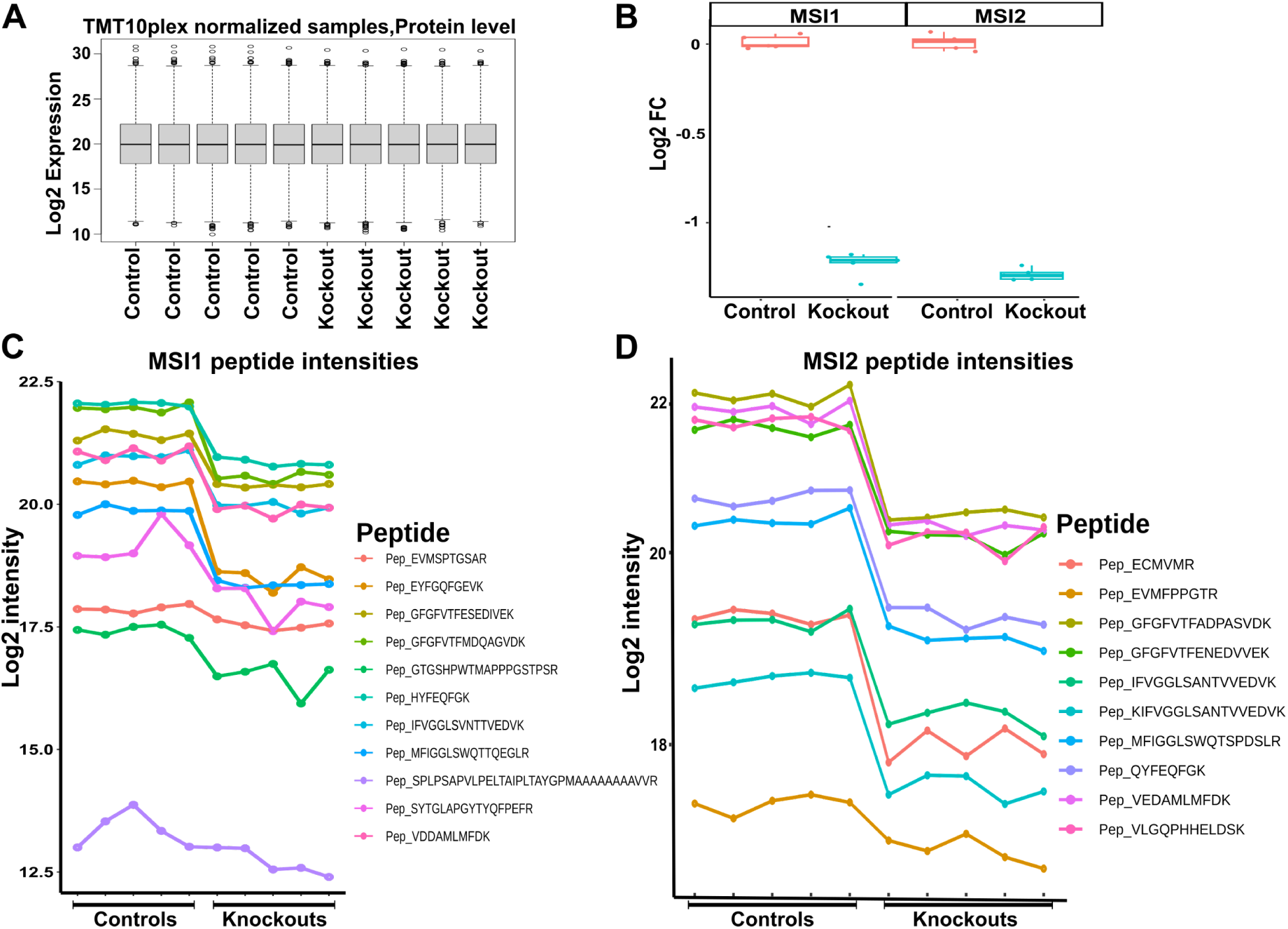
Decrease of MSI1 and MSI2 protein levels in the retina after induced double knockout of *Msi1* and *Msi2* in mature photoreceptor cells. **A)** Box plot showing the distribution of normalized signal intensities across samples analyzed by isobaric labeling and tandem MS (MS3). **B)** Box plots showing the log2 of the fold difference of MSI1 and MSI2 protein levels in the retina of control and knockout mice relative to the median of the controls. Changes in the levels of individual tryptic peptides identified for MSI1 **(C)** and MSI2 **(D)** in control and knockout retina.

**Figure 7 - figure supplement 2.**
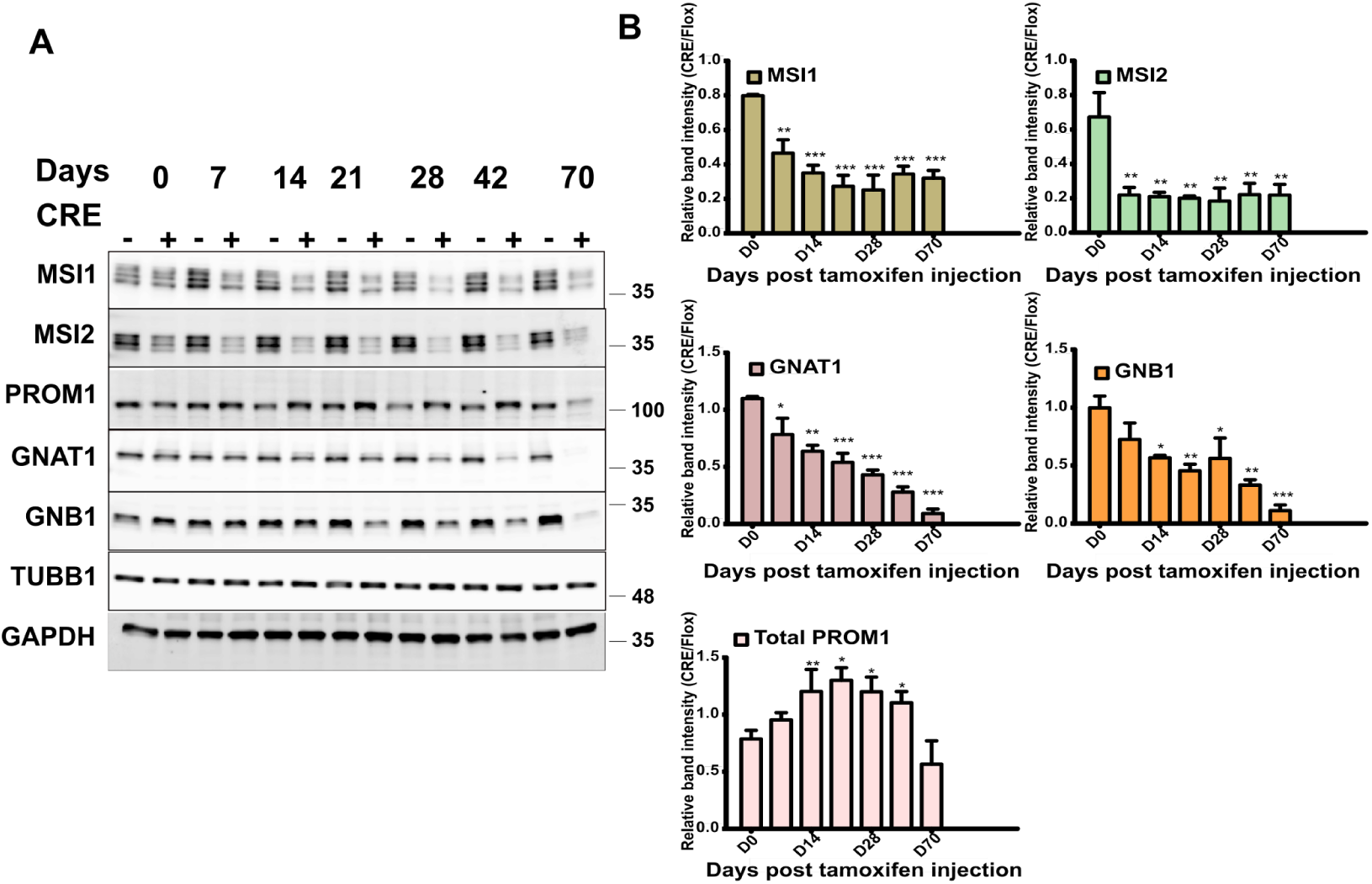
Validation of MS3 data by western blot. **A)** Representative immunoblot showing levels of selected proteins after combined deletion of *Msi1* and *Msi2* in mature photoreceptors. B) Quantification of western blot data for MSI1, MSI2, GNAT1, GNB1 and PROM1. TUBB1 and GAPDH were used as controls to normalize for loading. Error bars represent standard error of the mean (SEM, n=3). The statistical significance of the pairwise comparisons of the protein levels at different time points to the baseline level at the day of the tamoxifen injection is indicated as: * p-value < 0.05, ** p-value < 0.01, *** p-value < 0.001.

